# Cytokinins regulate spatially-specific ethylene production to control root growth in *Arabidopsis*

**DOI:** 10.1101/2023.01.07.522790

**Authors:** Amel Yamoune, Marketa Zdarska, Thomas Depaepe, Anna Korytarova, Jan Skalak, Kenneth Wayne Berendzen, Virtudes Mira-Rodado, Paul Tarr, Eliska Spackova, Lucia Badurova, Barbora Parizkova, Abigail Franczyk, Ingrid Kovacova, Marketa Pernisova, Ondrej Novak, Elliot Meyerowitz, Klaus Harter, Dominique Van Der Straeten, Jan Hejatko

## Abstract

The two principal growth regulators cytokinins and ethylene are known to interact in the regulation of plant growth. However, information about underlying molecular mechanism and positional specificity of the cytokinin/ethylene crosstalk in root growth control is scarce. We have identified spatial specificity of cytokinin-regulated root elongation and root apical meristem (RAM) size, both of which we demonstrate to be ethylene biosynthesis-dependent. Upregulation of the cytokinin biosynthetic gene *ISOPENTENYLTRANSFERASE* (*IPT*) in proximal and peripheral tissues leads to both root and RAM shortening. In contrast, *IPT* activation in distal and inner tissues reduces RAM size while leaving the root length comparable to mock-treated controls. We show that cytokinins regulate two steps specific to ethylene biosynthesis, the production of ACC by ACC SYNTHASEs (ACSs), and its conversion to ethylene by ACC OXIDASEs (ACOs). We describe cytokinin- and ethylene-specific regulation controlling the activity of *ACSs* and *ACOs* that are spatially discrete along both proximo/distal and radial root axes. Using direct ethylene measurements, we identify *ACO2, ACO3* and *ACO4* as being responsible for ethylene biosynthesis and the ethylene-regulated root and RAM shortening in cytokinin-treated *Arabidopsis*. Finally, we describe the tight cooperation between cytokinin and ethylene signaling in cytokinin-induced, ethylene-regulated control of *ACO4* due to the direct interaction between ARABIDOPSIS RESPONSE REGULATOR 2 (ARR2), a member of the multistep phosphorelay cascade and the C-terminal portion of ETHYLENE INSENSITIVE 2 (EIN2-C), a key regulator of canonical ethylene signaling.

## Introduction

Roots or root-like structures are one of the key adaptations in plants that were critical for terrestrial colonization (Hetherington and Dolan, 2018). Roots mediate a number of biotic and abiotic interactions (Bakker et al., 2018; Comas et al., 2013) and root architecture is one of the key yield-determining traits under both normal and stress (particularly drought) conditions (Lynch, 2007; Ramireddy et al., 2018; Uga et al., 2013). Understanding the factors controlling root growth is critical for building a comprehensive picture of plant developmental and adaptive responses that directly impact crop productivity.

The overall growth rate in the root is determined by the balance between three fundamental processes – i) cell proliferative activity in the root apical meristem (RAM), ii) cell differentiation and iii) elongation of cells leaving the RAM. All of these processes are known to be under the control of phytohormones including cytokinins and ethylene [reviewed by (Kong et al., 2018; Svolacchia et al., 2020; Takatsuka and Umeda, 2014; Yamoune et al., 2021)]. Cytokinins control RAM size and its proliferation capacity both in a positive and negative way. Cytokinins increase RAM size by enhancing stem cell proliferation, but can also shorten the RAM (a process involving crosstalk with auxin and gibberellic acid) by inducing cell differentiation in the root transition zone [for a recent review see (Svolacchia et al., 2020; Yamoune et al., 2021)]. The involvement of cytokinin-regulated auxin transport has been invoked in the regulation of root cell elongation both in an ethylene-dependent and -independent manner (Street et al., 2016), possibly by inducing cell wall stiffening (Liu et al., 2022).

Ethylene is one of the main regulators of root cell elongation, with an inhibitory effect that has been known for decades (Dolan, 1997). This ethylene-mediated inhibition of cell elongation is not limited to the root, and is to a large extent, if not exclusively, dependent on ethylene-regulated auxin biosynthesis and transport (Hu et al., 2017; Mazzoni-Putman et al., 2021; Stepanova and Alonso, 2009; Vaseva et al., 2018; Zemlyanskaya et al., 2018). Continuous treatment with the ethylene biosynthesis precursor 1-aminocyclopropane-1-carboxylate (ACC) leads to the inhibition of cell elongation by repressing cell elongation-promoting factors as well as inducing genes whose products attenuate cell elongation (Markakis et al., 2012). Despite this, the early ethylene response can be both positive and negative (depending on the developmental context and its position in the RAM epidermis) and this effect seems to be independent of the role of ethylene in inducing cell differentiation (Le et al., 2001). Apart from its role in cell elongation, ethylene has also been demonstrated to control cell division in the root stem cell niche (Ortega-Martinez et al., 2007) and participates (along with cytokinins) in the control of RAM size by inducing cell differentiation in the root transition zone (Street et al., 2015; Zdarska et al., 2019).

Ethylene biosynthesis in plants starts by the conversion of methionine by S-adenosyl-L methionine (SAM) synthetase to S-adenosyl methionine (SAM), the general ethylene precursor shared by several metabolic pathways. SAM serves as a substrate for ACC SYNTHASEs (ACSs), mediating the first (and rate-limiting) step dedicated exclusively to ethylene biosynthesis, leading to the formation of 1-aminocyclopropane-1-carboxylic acid (ACC). ACC oxidation to ethylene by ACC OXIDASEs (ACOs) is the second and final step specific to the ethylene biosynthetic pathway. Given the key importance of ethylene in many aspects of the plant life cycle, it is not surprising that the activity of both ACSs and ACOs are under tight transcriptional and posttranscriptional control. Moreover, the levels of the non-proteinogenic amino acid ACC can be further regulated by conjugation and translocation. For more detailed information on ethylene biosynthesis see recent reviews by (Depaepe and Van Der Straeten, 2020; Pattyn et al., 2021).

Ethylene is perceived by the ethylene-responsive sensor histidine kinases ETHYLENE RESPONSE 1 (ETR1) and ETHYLENE RESPONSE SENSOR 1 (ERS1) and by the HK-like Ser/Thr kinases ETR2, ERS2 and ETHYLENE INSENSITIVE 4 (EIN4) [reviewed in (Binder, 2020; Etheridge et al., 2006; Chen et al., 2005)]. The downstream target of ER-located ethylene sensors in the canonical ethylene signaling pathway is the Raf family Ser/Thr kinase CONSTITUTIVE TRIPLE RESPONSE 1 (CTR1) (Kieber et al., 1993). Both the receptors and CTR1 act as negative regulators of the signaling pathway. Ethylene binding switches off the ethylene sensors, attenuating the CTR1-mediated phosphorylation of the ER-associated N-ramp like protein EIN2. As a result, the C-terminal end of hypo-phosphorylated EIN2 (EIN2-C) is disinhibited. In the cytoplasm, EIN2-C initiates degradation of the mRNA of *EIN3-BINDING F BOX PROTEIN* (*EBF1*) and *EBF2*, leading to the stabilization of the ethylene-responsive transcription factor EIN3. In parallel, EIN2-C translocates into the nucleus, becoming part of the complex facilitating EIN3-regulated transcription (Binder, 2020; Ju and Chang, 2012; Li et al., 2015; Wen et al., 2012).

Cytokinin signaling is also initiated by histidine kinases, but the downstream response, unlike for ethylene, is mediated via a multistep phophorelay (MSP) pathway, also called two-component signaling [for a review see (Kieber and Schaller, 2018; Leuendorf and Schmuelling, 2021; Mira-Rodado, 2019)]. In the MSP pathway, cytokinins are perceived by the CHASE domain of ARABIDOPSIS HISTIDINE KINASE 2 (AHK2), AHK3 and AHK4, leading to the autophosphorylation of a conserved His. That triggers the His-to-Asp-to-His-to-Asp downstream phosphorelay and activation (via phosphorylation of their conserved Asp residue) of nuclear-localized type B ARABIDOPSIS RESPONSE REGULATORs [RRBs (Heyl et al., 2013)], acting as cytokinin-regulated transcription factors.

Regulation of root growth involves tight cytokinin/ethylene crosstalk. Exogenous cytokinins have an inhibitory effect on root cell elongation (Beemster and Baskin, 2000) that is mediated via cytokinin-induced ethylene production; the inhibitory effect of cytokinins on the elongation of both root and hypocotyl cells was shown to be dependent on functioning ethylene signal transduction (Cary et al., 1995; Ruzicka et al., 2009). In line with that, the regulatory effect of cytokinins on both RAM size and root cell elongation in rice were demonstrated to be mediated by increased ethylene content (Zou et al., 2018). Mechanistically, cytokinin and ethylene interact at the level of both biosynthesis as well as signaling. Tight interaction between MSP and canonical ethylene signaling has been reported [for a recent review see (Skalak et al., 2021)]. Briefly, ETR1 was shown to mediate ethylene-regulated MSP signaling in the root transition zone to control RAM size via ethylene-induced cell differentiation (Street et al., 2015; Zdarska et al., 2019). The action of ETR1 was proposed be mediated via ETR1-induced phosphorylation of the histine kinase AHK5 (Szmitkowska et al., 2021), eventually leading to the phosphorylation of RRB ARR2 (Hass et al., 2004). In rice, the ethylene sensor OsERS2 was shown to interact with the AHK5 orthologue MHZ1/OsHK1 and control its HK activity (Zhao et al., 2020). Cytokinins were also demonstrated to upregulate ethylene biosynthesis by stabilizing ACS5 and ACS9 (Hansen et al., 2009; Chae et al., 2003; Rashotte et al., 2005; Vogel et al., 1998).

Here we describe the identification of a tight interaction network between cytokinins and ethylene biosynthetic genes. We show that cytokinins not only stimulate ACC production by transcriptional regulation of several ACSs, but that they also regulate the last step of ethylene biosynthesis by activating ACOs. We show that cytokinins control root elongation and RAM size by inducing ethylene biosynthesis in a spatially distinct manner and we describe a novel mechanism of interaction between MSP and canonical ethylene signaling driving the expression of *ACO4*.

## Results

### Cell type-specific cytokinin overproduction is necessary to induce ACC biosynthesis and root shortening

Exogenously applied cytokinins inhibit root cell elongation primarily through cytokinin-induced ethylene production: cytokinin-induced root shortening depends on functional ethylene signaling or ethylene biosynthesis [(Ruzicka et al., 2009) and Supplemental Figure 1]. To assess the potential cell-type specificity of cytokinin-induced root shortening, we upregulated cytokinin biosynthesis in the outer RAM cell layers (epidermis and cortex), shown to be required for ethylene-regulated root growth (Vaseva et al., 2018), and in the more internal (provascular/stele) tissues, suggested to be important for cytokinin-mediated RAM shortening (Dello Ioio et al., 2007). This was achieved by activating the cytokinin biosynthetic gene *ISOPENTENYLTRANSFERASE* (*IPT*) in a cell type-specific manner using the GAL4>>UAS activator-reporter system (Bielach et al., 2012; Laplaze et al., 2005). *IPT* upregulation in the epidermis/cortex of the root transition/elongation zone and in more proximal differentiated tissues in J2601>>*IPT* is associated with a strong inhibition of root growth and a stimulation of root hair formation (Figures 1A, B), both of which were not observed in the presence of 2-aminoethoxyvinylglycine (AVG), an inhibitor of ACC biosynthesis (Yang and Hoffman, 1984). In contrast, *IPT* activation in provascular tissues in J2351>>*IPT* resulted in no root length change and a weaker induction of hairy root formation compared to epidermis/cortex stimulated *IPT*, although the roots were still sensitive to exogenously added ACC (Figures 1A, B). Nonetheless, significant RAM reduction was observed upon *IPT* activation in both J2601>>*IPT* and J2351>>*IPT* (Figures 1C, D). Upregulation of *IPT* expression in both epidermis/cortex and provascular/stele tissues resulted in increased endogenous ACC levels. However, the cytokinin-induced ACC upregulation was more prominent in J2601>>*IPT* (4.3-fold change) than in J2351>>*IPT* (1.6-fold change; Figure 1E).

**Figure 1.**
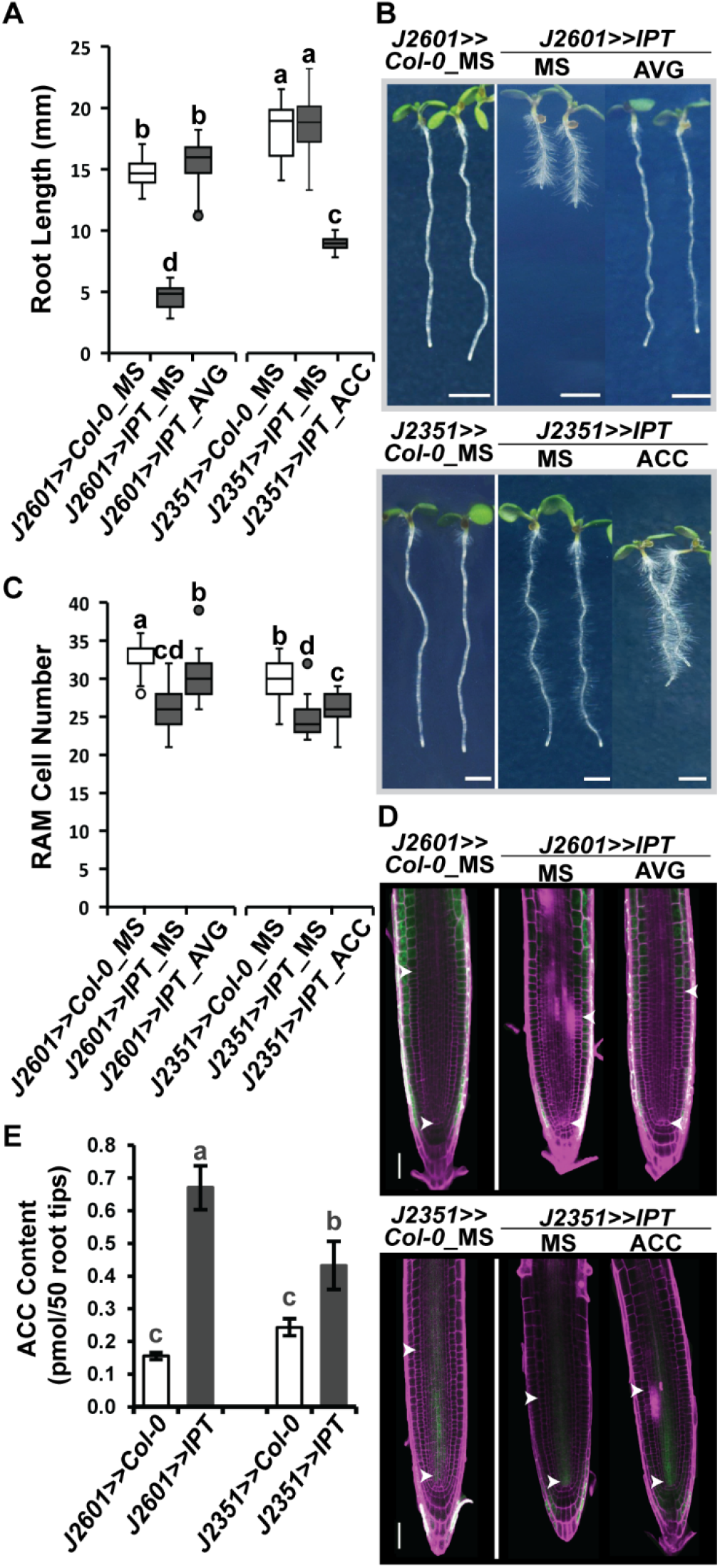
IPT ectopic overexpression in the epidermis induces root shortening. Root length, (**B**) RAM cortex cell number and **(C, D)** the representative images of six-day-old seedlings of *IPT* overexpressing lines: *J2601>>IPT* (epidermis/cortex) grown ½MS media with or without 0.2 µM AVG and *J2351>>IPT* (stele/LRC) grown on ½MS media with or without 1 µM ACC; untreated *J2601>>Col-0* and *J2351>>Col-0* were used as controls, respectively. (**E**) ACC level in root tips of *J2601>>IPT* and *J2351>>IPT* lines and their respective controls *J2601>>Col0* and *J2351>>Col0*. The boxplots in (A) and (B) represent data from three independent replicas and the bars in (E) the means ± SD of three biological replicates; the letters represent significance classes determined by a linear mixed model ANOVA and Tukey’s post hoc HSD test. The white arrowheads in (D) mark the extent of the RAM. The scale bars represent 2.5 mm in and 100 µm in (D).

These results confirm the tight interaction between cytokinins and ethylene biosynthesis in controlling root growth, and show that cell type-specific induction of cytokinin biosynthesis seems to be important for cytokinin-induced ACC production and root shortening. While cytokinin upregulation in both distal/internal and proximal/outer tissues leads to RAM shortening (in former case without a significant effect on root elongation), the activation of endogenous cytokinin production in the proximal/outer cell types is necessary for the cytokinin-induced, ethylene-mediated root growth reduction.

### Both cytokinin and ethylene upregulate transcription of *ACC SYNTHASES*

To identify the molecular events mediating cytokinin-induced ACC synthesis in *Arabidopsis* roots, we assayed the cytokinin response of transcriptional *pACS::GUS* reporters (Tsuchisaka and Theologis, 2004) as well as newly prepared lines carrying *ACS2* (*pACS2::ACS2:GFP*) and *ACS7* (*pACS7::ACS7:GFP*) translational fusions (Figure 2A, B, Supplemental Figure 2). Of the eight investigated *ACS* genes, the activities of five were induced by cytokinin treatments. The activities of *ACS5, ACS6, ACS8* and *ACS11* were enhanced in most cell types in the differentiation/elongation zone as well as in older parts of the root. Cytokinin treatment strongly upregulated *ACS5, ACS6* and *ACS8* activity also in the vasculature and stele of the root tip while *ACS7* activity was induced specifically in the epidermal and cortical cells of the root transition zone. The remaining genes were cytokinin-insensitive (*ACS2* and *ACS4)* or were activated only very weakly (*ACS9*; Supplemental Figure 2). *ACS7* was alone among the cytokinin-induced ACSs in responding specifically to cytokinins, as revealed by comparing cytokinin treatment with and without AVG (Figures 2A, B). *ACS5, ACS6, ACS8* and *ACS11* revealed a combination of both cytokinin-induced and cytokinin-induced, ethylene-mediated activation (i.e., activation dependent on ACC production), often being spatially restricted mostly to the stele/vasculature and non-vascular cell types located proximal to the root transition zone (Figure 2A). In line with the observed cytokinin- and ethylene-responsiveness of several *ACS* genes, we found a reduced sensitivity to cytokinin-induced RAM shortening particularly in *acs6, acs7* and *acs8* single and *acs2 acs6* and *acs5 acs9* double mutant *Arabidopsis* lines. Moreover, smaller RAMs were observed also under control conditions in single *acs5, acs6, acs7* and *acs9* mutant lines when compared to WT Col-0 (Figure 2C).

**Figure 2.**
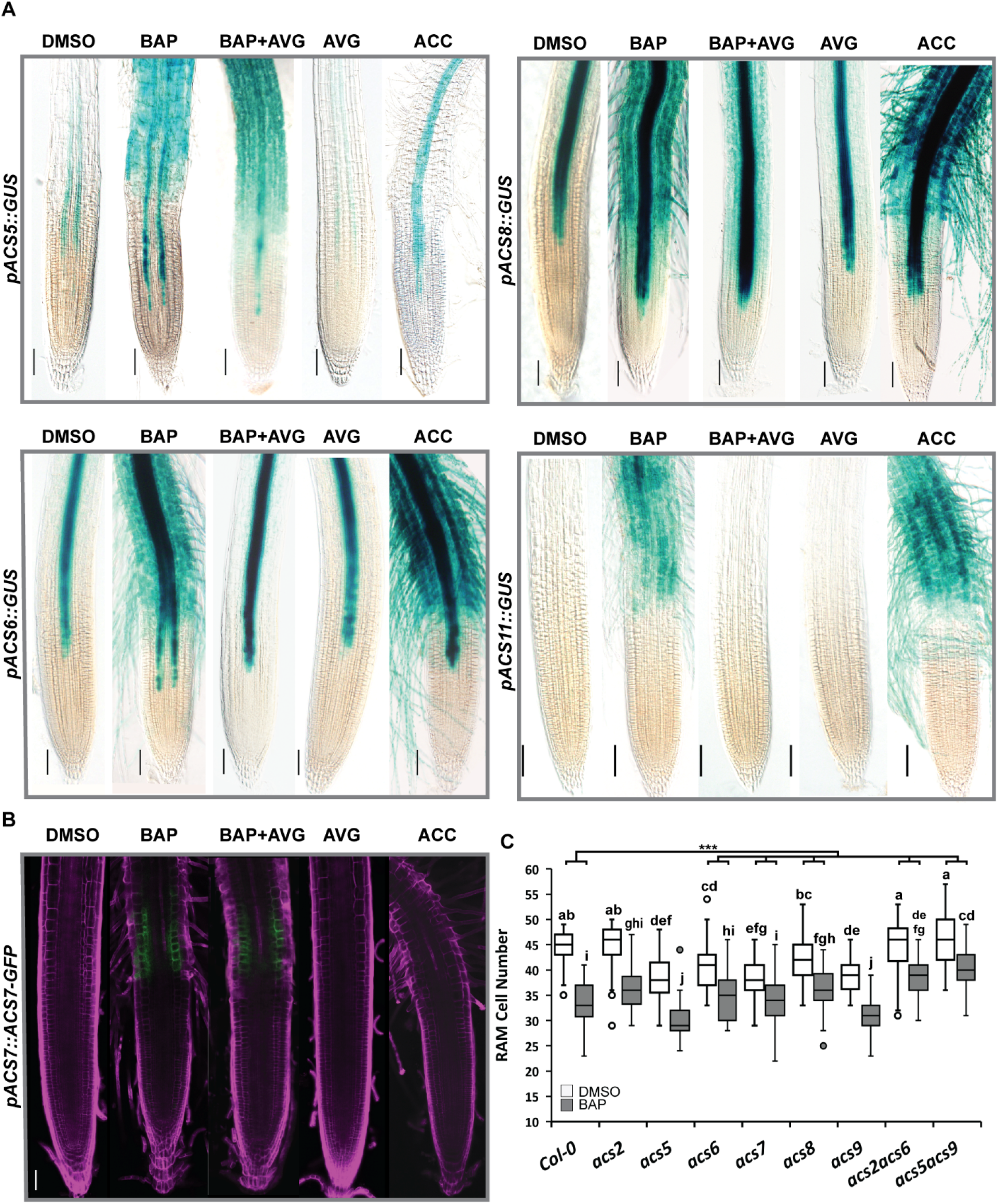
Cytokinin induces the expression of several *ACC SYNTHASE* genes. (**A**) Six-day-old seedlings of *pACSx*::GUS (*ACS5, ACS6, ACS8* and, *ACS11*) transcriptional reporter and (**B**) *pACS7*::ACS7:GFP translational fusion lines exposed for 24h to different hormones (5 µM BAP, 5 µM BAP + 1 µM AVG, 1 µM AVG, 5 µM ACC; control is 0.01% DMSO) in liquid media. Scale bars represent 100 µm. (**C**) RAM cortex cell number of six-day-old *WT Col-0* and *acs* mutant lines (*acs2, acs5, acs6, acs7, acs8, acs9, acs2acs6* and *acs5acs9)* treated for 24h with 5 µM BAP (control is 0.01% DMSO). Boxplots represent data from the three independent replicas, the letters show significance classes determined by a linear mixed model ANOVA followed by Tukey’s post hoc HSD test (see supplemental methods). The line-tree at the top of the graph in (C) represents the difference-in-differences (DD) estimation between the BAP-reduced RAM size change in *WT Col-0* compared to the change in the different *acs* knockouts; the asterisks denote significance at p<0.001. The scale bars represent 100 µm in (A) and 50 µm in (B).

Taken together, our findings imply that cytokinins upregulate ACC production in *Arabidopsis* root through the cell type-specific transcriptional regulation of several *ACS* genes employing both cytokinin-specific as well as ethylene-dependent mechanisms. Cytokinin-inducible *ACS5, ACS6, ACS7* and *ACS9* regulate RAM size under control conditions and, together with *ACS8*, are necessary for cytokinin-induced RAM shortening.

### ACO2, ACO3 and ACO4 are controlled by cytokinins and cytokinin-induced ethylene

Our previous findings revealed a possible role for cytokinins as positive regulators of *ACOs* (Zdarska et al., 2013). Accordingly, we observed that α-aminoisobutyric acid (AIB), an inhibitor of ACO activity (Satoh and Esashi, 1982; Satoh and Esashi, 1983), partially rescued cytokinin-induced root shortening (Supplemental Figures 1A, B), suggesting a possible role for *ACOs* in cytokinin-regulated root growth. Using RT-qPCR and/or newly-prepared reporter lines, we found that exogenously applied cytokinin significantly upregulates *ACO3* and *ACO4*, but downregulates *ACO1, ACO2* and *ACO5* in the root tip (Figures 3A, B; Supplemental Figure 3). A contrasting effect of cytokinins and ethylene/ACC regulation was observed in case of *ACO2*. In the epidermis of the transition zone/elongation zone, *ACO2* was downregulated by cytokinins, but upregulated by ACC (Supplemental Figures 3A, B). Furthermore, cytokinin-induced upregulation of *ACO2* and *ACO3* was observed in the vasculature of the fully differentiated proximal portion of the root (Supplemental Figures 3I, J). As with ACSs, we observed a combinatorial effect of both cytokinin- and ethylene-specific regulation also for *ACO3* and *ACO4*. Cytokinin-specific activation of *ACO3* was detected in the stele and the vasculature of the root transition/elongation zone. The strong *ACO3* activation in the vascular tissues of the more proximal portion of the root (early differentiation zone) turned out to be mediated via cytokinin-induced ethylene production (Figure 3C, Supplemental Figures 3C-E). *ACO4* was upregulated in the columella and lateral root cap (LRC) in both a cytokinin- and an ethylene-specific manner, while only ethylene-specific activation was observed in the epidermis of the root transition/early elongation zones (Figure 3D, Supplemental Figures 3F-H).

**Figure 3.**
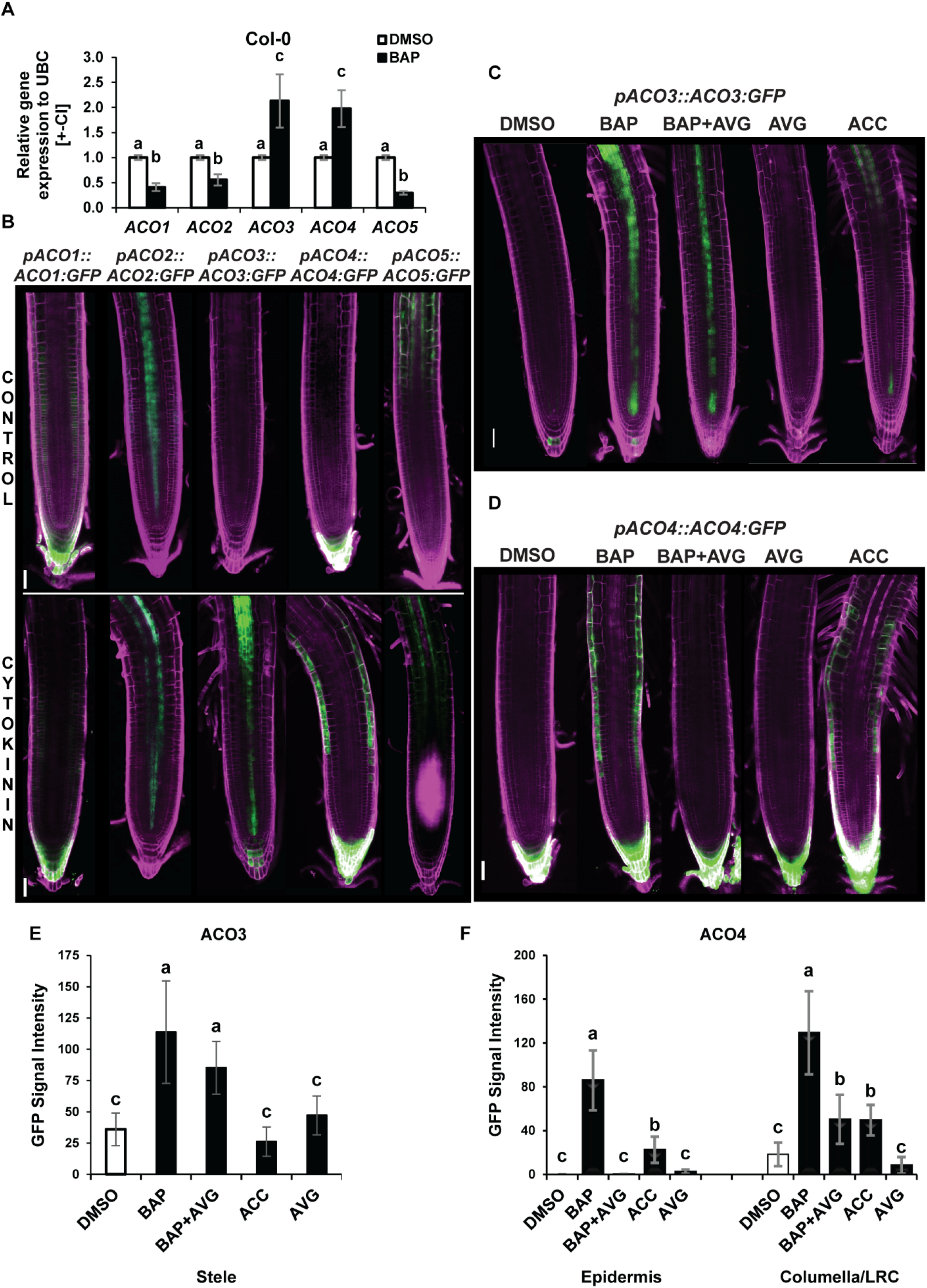
Hormonal control over *ACO3* and *ACO4* genes. RT qPCR quantification of the *ACOs* (*ACO1-5*) transcripts in six-day-old *WT Col-0* root tips following 6h of treatment with 5 µM BAP; 0.01% DMSO control. Relative gene expression is normalized to *UBC10* with mean values +/-CI of 4 biological replicates shown; the letters indicate statistically homogenous groups following the Kruskal-Wallis and Dunn post-hoc tests. Root tips of six-day-old ACO translation fusions (*pACO1-5::ACO1-5:GFP*) treated for 24 h with 5 µM BAP; 0.01% DMSO control. (**C**) Hormonal control of ACO3 and (**D**) ACO4 and (E, F) corresponding GFP intensities in the specified tissue files visualized in the root tip of six-old-old seedlings *pACO3::ACO3:GFP* and *pACO4::ACO4:GFP*, respectively, treated for 24h with 5 µM BAP, 5 µM BAP + 1 µM AVG, 1 µM AVG or 5 µM ACC; control is 0.01% DMSO. The areas used for GFP intensity measurements are shown in Supplemental figure 3. The scale bars (B-C) represent 50 µm.

In conclusion, besides inducing ACC production, cytokinins are also spatially specific regulators of ACC oxidation, the last step in ethylene biosynthesis. As for cytokinin-activated *ACSs*, cytokinins control *ACOs* both directly as well as via cytokinin-induced ethylene production.

### Multistep phosphorelay and canonical ethylene signaling are necessary for, and cooperate in, cytokinin-induced upregulation of ACO3 and ACO4

To find the molecular mechanism underlying the cytokinin-induced upregulation of *ACO3* and *ACO4, pACO3::ACO3-GFP* and *pACO4::ACO4-GFP* were introduced by crossing into various mutant backgrounds deficient in multistep phosphorelay (*arr1-3, arr2-5, arr10*-*1* and *arr12*-*1*) and/or the canonical ethylene signaling (*ein2-1*). We found that ARR1 was necessary for cytokinin-mediated upregulation of *ACO3*, whereas both functional ARR2 and EIN2 were required for ethylene-dependent activation of *ACO4* (Figures 4A, B). To get a more detailed mechanistic insight into *ACO* regulation, we assayed the ability of *ACO3* and *ACO4* promoters to physically interact with the RRBs and EIN2-C in a yeast one-hybrid (Y1H) assay. The N-terminal receiver domain of RRBs was previously demonstrated to act as a phosphorylation-dependent negative regulator of RRBs binding to DNA (Sakai et al., 2000a). Therefore, the interaction of *ACO3/4* promoter fragments was tested with truncated RRB versions (ΔDDKARR1, ΔDDKARR2, ΔDDKARR10, and ΔDDKARR12, Figure 4C, Supplemental Figure 4A), consisting only of the GARP DNA-binding domain. Among the tested TFs, only ΔDDKARR1 was able to bind fragments of the *ACO3* promoter (Figure 4D). *ARR1* was inducible by both cytokinins and ethylene (Supplemental Figure 4B). Neither ΔDDKARR2 nor EIN2-C were able to bind the *ACO4* promoter when expressed separately. However, when co-expressed, ΔDDKARR2 and EIN2-C allowed the activation of the yeast reporter that was under the control of *ACO4* promoter fragments (Figure 4E), suggesting their cooperative binding to *pACO4*. As EIN2-C does not possess a DNA-binding domain (Zhang et al., 2016), we presume that the ARR2 (ΔDDKARR2) GARP domain might mediate the recruitment of an ARR2/EIN2-C complex to the *ACO4* promoter. Accordingly, FLIM-FRET detected a strong interaction between ΔDDKARR2 and EIN2-C transiently produced in tobacco leaves (Figures 4F, G).

**Figure 4.**
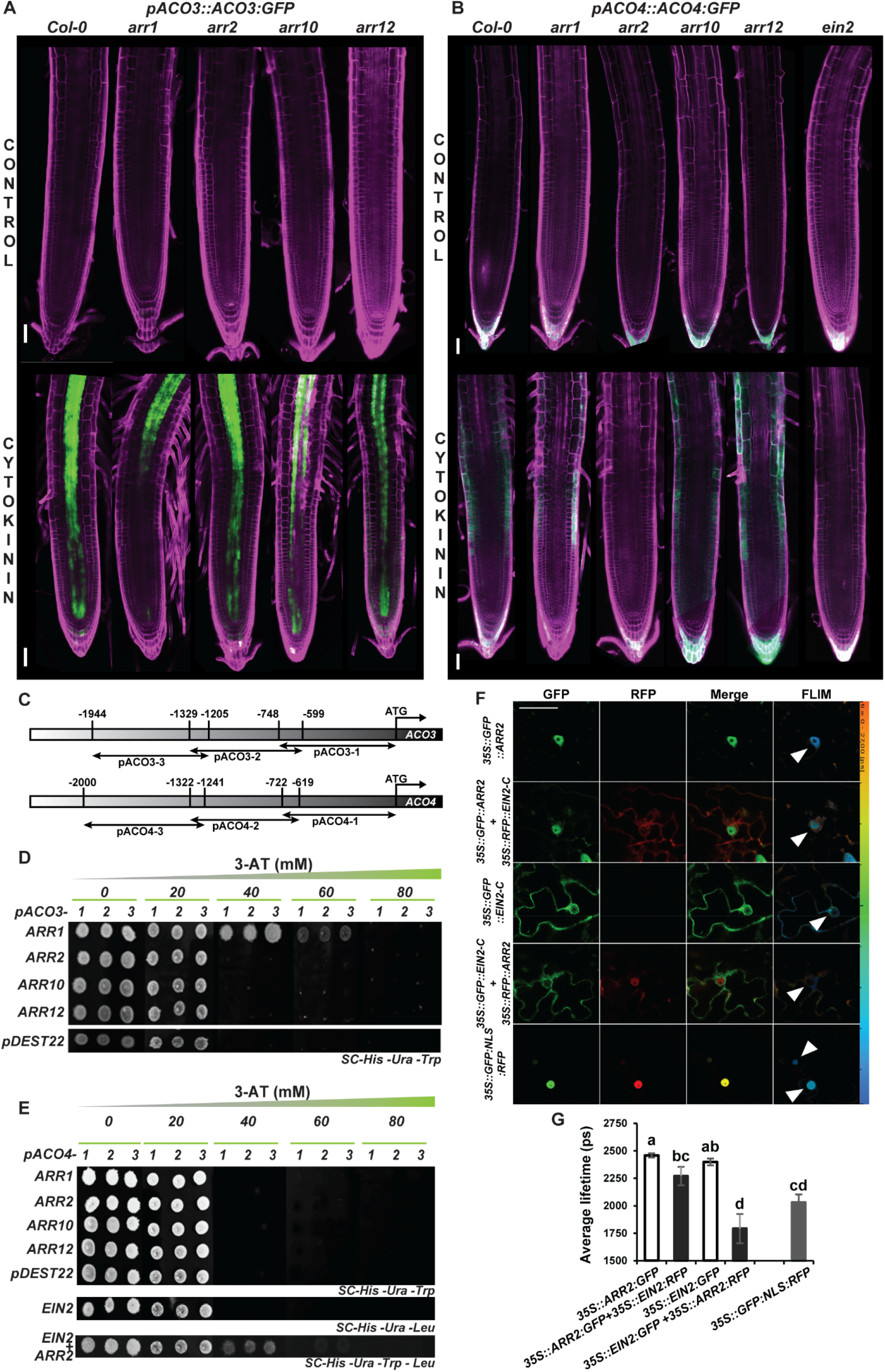
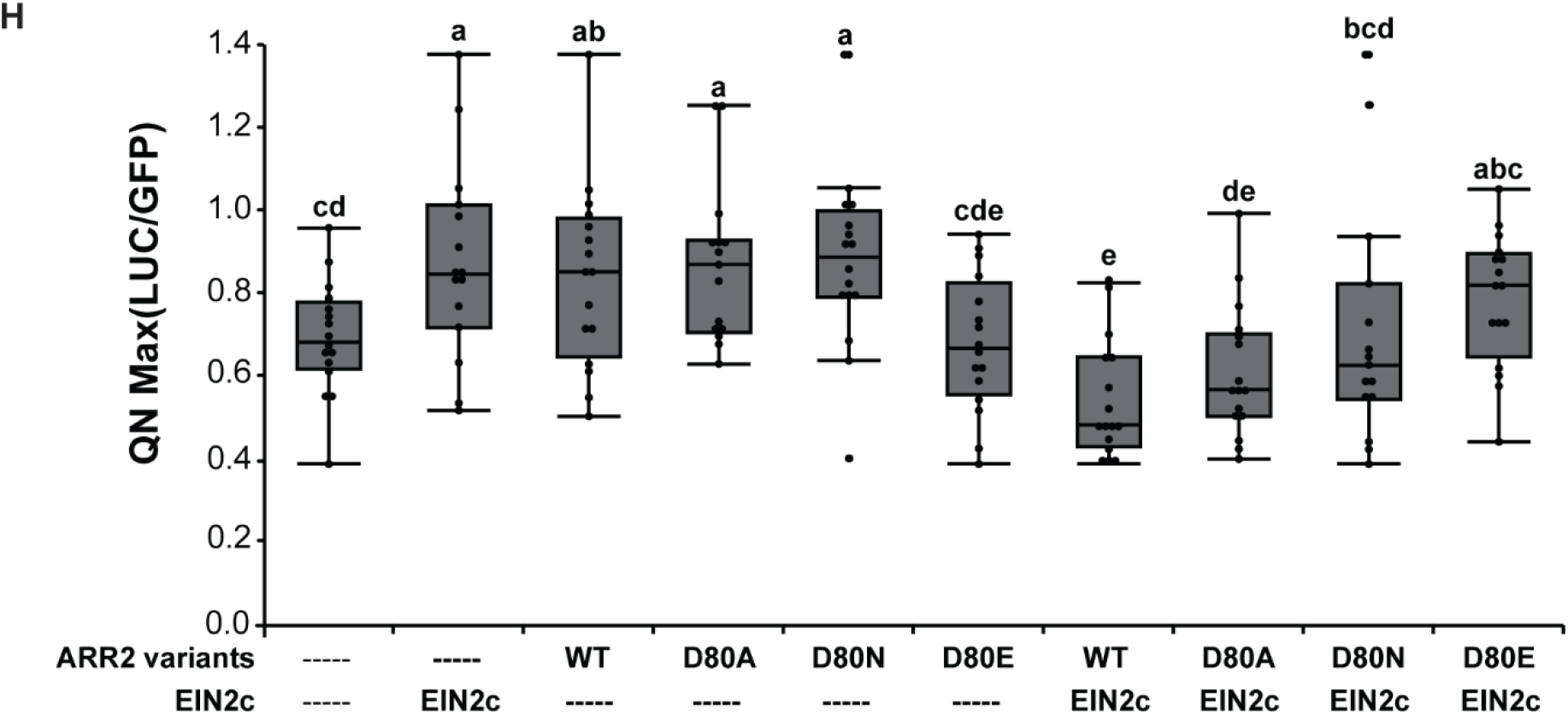
ARR1 directly binds *ACO3* whereas both ARR2 and EIN2 associate to initiate the transcription of *ACO4*. (**A**) *pACO3::ACO3:GFP* and **(B)** *pACO4::ACO4:GFP* in *WT Col-0* and genetic backgrounds deficient in type-B *ARRs* or *EIN2* treated for 24h with 5 µM BAP; control is 0.01% DMSO. **(C)** Schematic representation of *ACO3* (upper) and *ACO4* (lower) promoter fragments (−1, -2, -3) used in the Y1H. (**D**) Y1H binding assay of truncated type-B ARRs TFs (ΔDDKARR1, ΔDDKARR2, ΔDDKARR10, ΔDDKARR12), and pDest22 (negative control) on *ACO3* as well as (**E**) truncated ARR-B, pDest22, and EIN2-C binding on *ACO4* promoter fragments; the interaction specificity was assayed in the presence of increasing concentrations of 3-Amino-1,2,4-triazole (3AT). (**F**) Representative confocal images and (**G**) the fluorescence lifetime measured in the FLIM-FRET interaction assay using the indicated vector combinations transiently expressed in *Nicotiana tabacum* leaves; *35S::GFP:NLS:RFP* was the positive control. The white arrowhead indicates FLIM measurement areas. Bars represent means ± SD of two biological replicates and letters indicate statistical significance (two-way ANOVA and Tukey’s post-hoc test). **(H)** Quantile normalization (QL) of LUC activity measured in living root-cell culture derived protoplasts transformed with p*ACO4::LUC* and the corresponding version of ARR2 and/or EIN2-C. A total of four experimental runs were merged by quantile normalization as in Wallmeroth *et* al. (2019). Boxplot and statistical tests were performed using JMP16. Significance classes were determined using Fisher’s least-squared test, α=0.05. D80A: non-phosphorylatable, fully inactive version of ARR2. D80N: non-phosphorylatable, partially inactive version of ARR2. D80E: phospho-mimic, constitutively active version of ARR2. The scale bars (A-B) represent 50 µm.

To further corroborate the functional importance of ARR2 and EIN2-C in the regulation of *ACO4* expression, we assayed *pACO4*-regulated gene expression in a LUC-based assay in protoplasts isolated from root suspension culture (Wallmeroth et al., 2019). When expressed separately, both EIN2-C and ARR2 were able to upregulate the activity of *pACO4*-controlled *LUC* (Figure 4H). While both WT and the two non-phosphorylatable versions of ARR2 (ARR2^D80A^ and ARR2 ^D80N^) activated *pACO4*, the dominantly-active ARR2^D80E^ (Hass et al., 2004) was unable to upregulate *pACO4* to levels above control. Furthermore, EIN2-C expressed together with both ARR2 WT and the non-phosphorylatable ARR2^D80A^ downregulated *pACO4*, but this negative effect could not be observed when EIN2-C was co-expressed with the dominantly-active ARR2^D80E^ (Figure 4H). We obtained comparable results for all constructs tested both in the presence and absence of 1 µM ACC (data not shown).

Taken together, our data indicate that both MSP and canonical ethylene signaling tightly cooperate to bring about the cytokinin-induced upregulation of ethylene biosynthetic genes in the root. Functional ARR1 is necessary for cytokinin-specific upregulation of *ACO3* in the stele and vasculature of the root transition/elongation zone. On the other hand, ARR2 and EIN2-C interact and mediate cytokinin-induced ethylene-dependent activation of *ACO4* in the root transition zone. The ability of the ARR2/EIN2-C complex to control *ACO4* expression seems to be dependent on the phosphorylation of ARR2 at its conserved Asp residue.

### ACO2, ACO3 and ACO4 are ethylene synthesizing enzymes involved in cytokinin-induced root and RAM shortening

To assess the functional importance of *ACOs* in cytokinin-induced ethylene biosynthesis and root growth, we measured ethylene formation in the roots of WT and several *aco* mutants, both in the presence and absence of cytokinins. Cytokinin treatment strongly upregulated ethylene production in WT. A statistically significant reduction in ethylene production compared to WT was detected in cytokinin-treated *aco2* single as well as *aco2 aco3* and *aco2 aco4* double mutants (Figure 5A). In the presence of cytokinins the *aco3* and *aco4* single mutant lines showed intermediate ethylene levels, statistically comparable to both WT and all ACO2-deficient lines.

**Figure 5.**
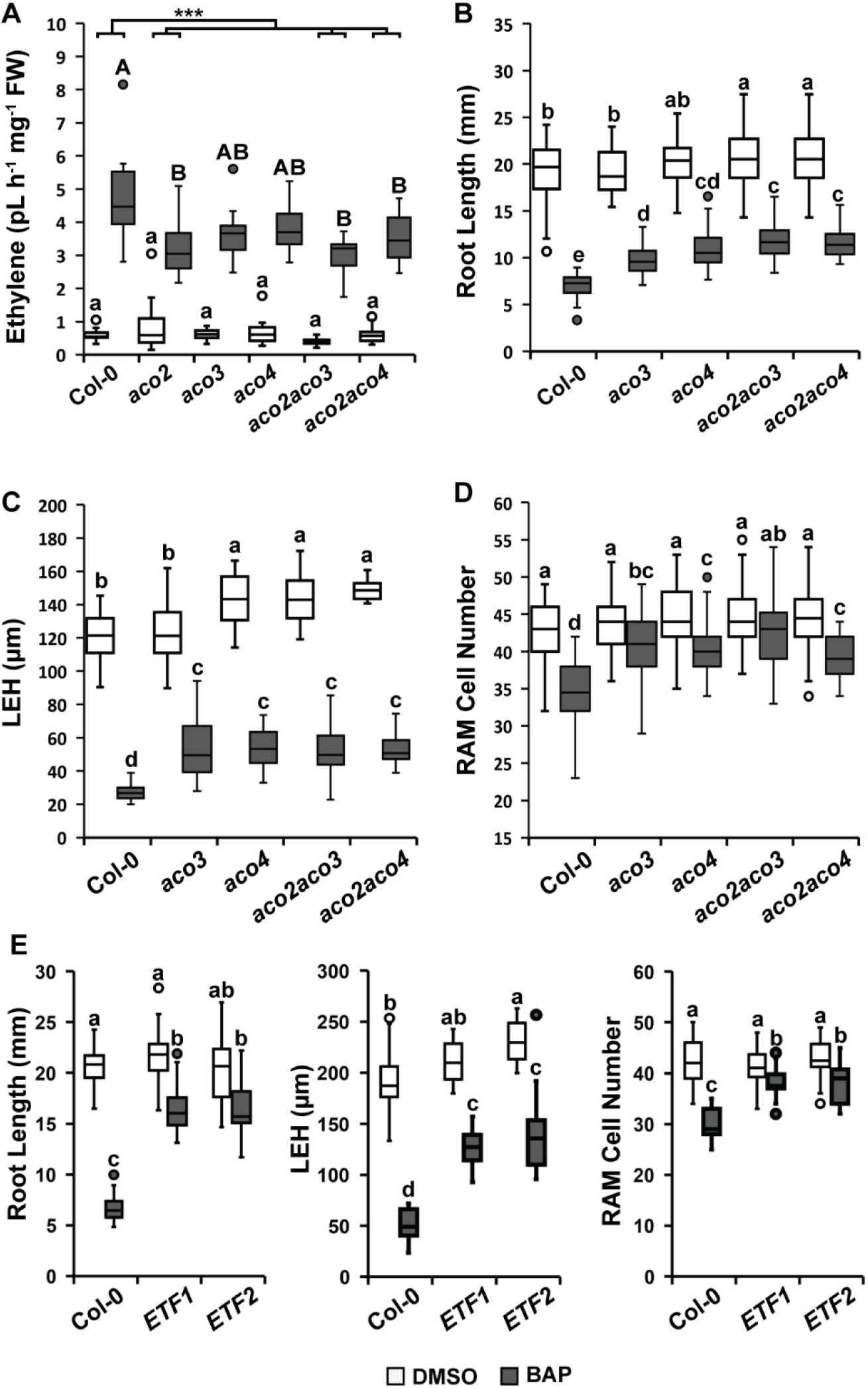
Cytokinin-induced ethylene impacts cytokinin-reduced RAM and root elongation. (**A**) Ethylene produced by the detached root (48 h of accumulation) of six-day-old *WT Col-0, aco2, aco3, aco4, aco2aco3* and *aco2aco4* seedlings treated with 5 µM BAP; control is 0.01% DMSO. (**B**) Root length, (**C**) the length of the first epidermal cell with a visible root hair bulge (LEH), and (**D**) RAM cortex cell number of six-day-old *WT Col-0, aco2, aco3, aco4, aco2aco3* and *aco2aco4* grown on +/-0.1 µM BAP ½MS; control is 0.01% DMSO. **(E)** The root growth parameters assayed in the presence and absence of cytokinins in ethylene-free (ETF) lines (Li et al., 2022). The boxplots represent data from three independent replicates, letters above the boxes represent statistically homogeneous groups after a linear mixed model ANOVA followed by Tukey’s post-hoc test (see supplemented methods). The black line-tree above A represents difference-in-differences (DD) estimation between BAP-treated *WT Col-0* and the different *aco* knockouts; the asterisks denote significance at p<0.001. The white boxes represent mock treated seedlings and the grey ones cytokinin treatments.

We then inspected the possible participation of *ACO*s in cytokinin-regulated root growth. Compared to WT, both *aco2aco3* and *aco2aco4* displayed longer roots under control conditions. A similar trend was also observable in the *aco4* single mutant line, although the difference was not statistically significant (Figure 5B). The inhibition of root length by cytokinin treatment was significantly lower in all tested *aco* mutants compared to WT. The largest drop in the sensitivity to cytokinin-mediated root shortening was observed in the *aco4* single and the *aco2aco3* and *aco2aco4* double mutants. A similar response was also seen upon measuring the length of the first epidermal cell showing a visible root hair bulge [LEH, (Le et al., 2001)]: all the tested mutant lines were less sensitive to cytokinins compared to WT. Importantly, the *aco4* single mutant as well as the *aco2aco3* and *aco2aco4* double mutant lines revealed extended LEH under control conditions, thus corresponding well with elongated roots (compare panels 5C and 5B). In case of RAM size, both *ACO3* deficient lines (*aco3* single and *aco2 aco3* double mutants) were either significantly less sensitive or completely resistant to cytokinin-induced RAM shortening, respectively. Though slightly weaker, a similar effect was observed for mutants deficient in *ACO4* (Figure 5D).

To further elaborate the possible contribution of remaining ACOs (ACO1 and ACO5) to cytokinin-regulated root growth, we determined the root growth parameters in the recently published ethylene-free lines (Li et al., 2022) deficient in all five assayed ACOs (ACO1-ACO5). Compared to all tested single and double *aco* mutants, we observed stronger decrease of the sensitivity in cytokinin-induced root and root cell shortening (Figure 5E). However, a similar level of insensitivity has been observed in cytokinin-induced RAM size in ethylene-free lines when compared to roots of ACO3-deficient single and double mutants (compare Figures 5D and E).

Altogether, our results demonstrate the involvement of *ACOs* in root growth either in the absence or presence of exogenous cytokinins. *ACO2, ACO3* and *ACO4*, and possibly also *ACO1* and/or *ACO5*, seem to contribute to ethylene-regulated cell elongation and RAM size. While *ACO3* plays a dominant role in ethylene regulation of RAM size, *ACO4* mediates ethylene control primarily through root cell elongation.

### Working model

Based on our data, we conclude that cytokinins regulate ethylene biosynthesis in a cell type-specific manner to control root growth at the level of both RAM activity and root cell elongation (Figure 6). Cytokinins are able to stimulate both ACC and ethylene production by inducing transcription of several *ACSs* and *ACOs* in a cytokinin-as well as an ethylene-specific fashion, which can be spatially traced to the stele/vasculature on the one hand and more peripheral tissue types (epidermis/cortex) on the other. Importantly, ethylene-specific regulation prevails in peripheral and proximal (transition zone and more proximal) tissues. Cytokinin-specific regulation, however, locates mostly to inner (vasculature/stele) and distal tissues, partially overlapping with the ethylene-specific regulation in proximal tissues, but also extending to the QC, as in the case of ACO3. This spatial specificity distinguishable in both longitudinal (proximodistal) and radial axes is important for the cytokinin-induced ethylene-mediated root shortening (taking place in peripheral and proximal tissues) and cytokinin-induced ethylene-dependent control of RAM size (prevailing in inner and distal cell types). In *cytokinin-induced ethylene-mediated root shortening* (orange box in Figure 6), the cytokinin-induced *ACS5* and *ACS5*/*7* in the stele and epidermis/cortex, respectively, of the root transition/elongation zone allow cytokinin-induced production of ACC that might be further metabolized by *ACO3*, which is itself induced by cytokinin-activated (possibly at the level of both transcription and phosphorylation) ARR1. The produced ethylene and/or ACC can further stimulate ACC production by upregulating *ACS6/8/9/11* and *ACO2* in the epidermis/cortex and/or *ACS5/6/8* in the vasculature and through the action of ARR2/EIN2C also *ACO4*. The ACO2/3/4-mediated ethylene may be a part of the feed-forward loop involved in ethylene-regulated root growth by attenuating the elongation of cells leaving the RAM. In the *cytokinin-regulated ethylene-dependent RAM size control* (green box in Figure 6), *ACS6, ACS8* and ARR1-regulated *ACO3* seem to mediate cytokinin-induced ACC/ethylene production more distally in the vasculature/stele of the transition zone/proliferation domain of the RAM that could be further potentiated by ACC-and/or ethylene-mediated *ACS6/8* and *ACO2* upregulation, eventually leading to the induction of cell differentiation and ethylene-dependent RAM shortening.

**Figure 6.**
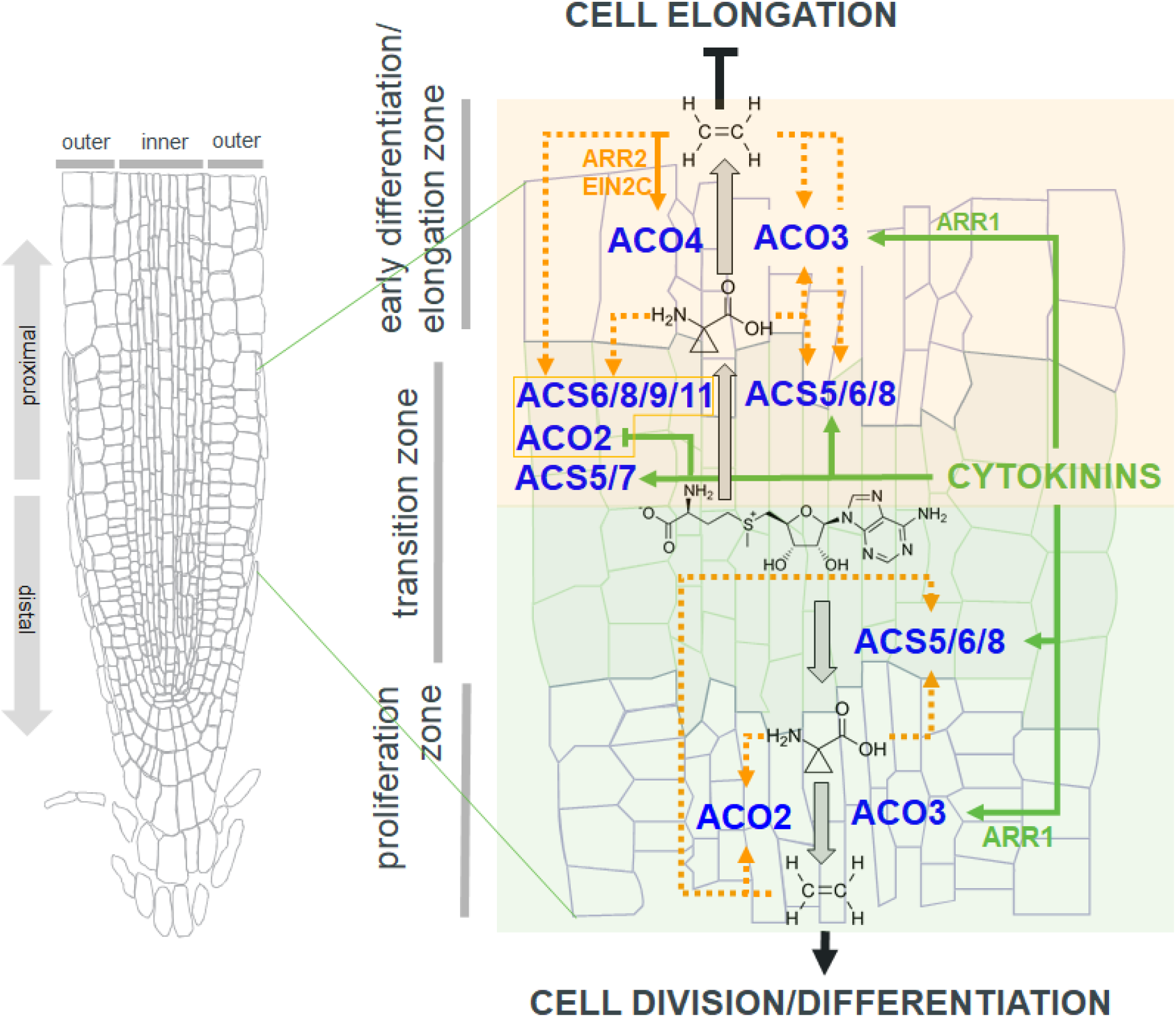
Working model. Cytokinins regulate spatial-specific expression of *ACSs* and *ACO* activity to control root growth. Cytokinin- and ethylene/ACC-specific regulations are in green and orange, respectively. Dotted lines are used wherever we cannot distinguish between ethylene- and ACC-mediated regulations. The *cytokinin-induced ethylene-mediated* regulation leading predominantly to root shortening via inhibition of cell elongation are highlighted by an orange box, the *cytokinin-regulated ethylene-dependent* regulation associated predominantly with RAM size control are highlighted by a green box. See the corresponding part of the main text for full descriptions.

## Discussion

### Cytokinins control root growth by regulating the expression of both *ACS* and *ACO*

So far, the only mechanism known to underlie cytokinin-induced ethylene production in *Arabidopsis* was the cytokinin-induced post-transcriptional stabilization of ACS proteins, specifically ACS2, ACS5, and ACS9 (Hansen et al., 2009; Chae et al., 2003; Lee et al., 2017; Vogel et al., 1998). The ACS stabilization does not appear to be specific to cytokinins, but can also be mediated by other hormones (Lee et al., 2017; Lee and Yoon, 2018). Nonetheless, this hormone-mediated ACS stabilization seemed thus far to be limited to etiolated seedlings (see the references above). Here we show that cytokinins upregulate *ACS* gene activity and transcript accumulation in light-grown *Arabidopsis* roots, as also demonstrated in rice, tobacco and tomato (Zhang et al., 2009; Zou et al., 2018). We further show that besides upregulating ACC production, cytokinins also control the last step of ethylene biosynthesis by differentially regulating the expression of *ACO* genes, controlling cell elongation and RAM size.

In *Arabidopsis, ACOs* are members of the large 2-oxoglutarate-dependent dioxygenase (2OGD) superfamily of non-heme iron-containing proteins that have diverse functions. Based on amino acid similarity, however, only five of the *ACOs* were proposed to be functional ACC oxidases in *Arabidopsis* [(Clouse and Carraro, 2014; Houben and Van de Poel, 2019; Sun et al., 2017) and references therein]. Recently, CRISPR/Cas-9-generated quintuple *aco1,2,3,4,5* mutant lines were obtained and the mutants were confirmed to be ethylene-free using gas chromatography (Li et al., 2022). This is in line with our direct ethylene measurements, suggesting that *ACO2, ACO3* and *ACO4* mediate ethylene biosynthesis in the root response to cytokinins. ACO2 seems to play a dominant role in the ethylene production in the entire root. This is in agreement with the previously identified non-transcriptional upregulation of ACO2 in the cytokinin-treated *Arabidopsis* root (Zd’arska et al., 2013) and possibly also reflects the fact that ACO2 is the most abundant ACO [Supplemental Figure 5; (Brady et al., 2007)], that is active throughout the differentiated root vasculature.

### Spatial and functional specificity of cytokinin-induced ethylene production

Our data show spatial specificity of cytokinin action in cytokinin-induced ethylene production and the consequent control of root growth. While increased production of cytokinins in the peripheral tissues located proximally to the root transition zone result in high ACC accumulation was associated with strong root size reduction, upregulation of cytokinin biosynthesis in the vasculature of the more distally located proliferation zone resulted in only a moderate ACC increase with a negligible effect on root length. This is in line with our findings, where most of the *ACS* genes were upregulated by cytokinins in either the transition zone of the root or more proximally, combining both cytokinin-specific and cytokinin-induced ethylene-mediated regulation. *ACO2* and *ACO4* seem to be important particularly in the latter, i.e., in a putative ethylene-mediated positive feedback loop upregulating several *ACSs*, and may act as a mechanism enhancing the effect of cytokinins on ACC and/or ethylene production in the elongating root cells. A similar mechanism (positive feedback regulation including ethylene-induced stabilization of ACS2 and ACS6) has been described under stress conditions (Vandenbussche et al., 2012). In contrast to this, there are fewer of ACSs under putative ACC- and/or ethylene-mediated positive feedback regulation in the distal RAM proliferation zone, possibly explaining the lower amount of ACC observed by *IPT* upregulation in the stele/vasculature-specific J2351 activator line. Nonetheless, even the (probably lower) amount of ethylene produced by ACO2 and ACO3 seems to be required for the RAM sensitivity to cytokinins, as clearly demonstrated by the nearly complete resistance of the *aco2 aco3* double mutant to cytokinin-induced RAM reduction. That was also observed at the level of ethylene signaling, where the ethylene-insensitive *etr1-1* as well as *etr1-9 ers1-3* complemented with HK-inactive ETR1 [ETR1-H/G2 (Hall et al., 2012)] was found to be resistant and/or less sensitive, respectively, to cytokinin-induced RAM shortening (Street et al., 2015; Zdarska et al., 2019). Here, we confirmed this phenomenon by identifying *ACO2/3* as being necessary for cytokinin-regulated RAM size.

However, the amount of cytokinin-induced ACC/ethylene may not be the only difference associated with position-specific cytokinin effects on the root growth. The spatial specificity that we observed for cytokinin-upregulated *ACSs* and *ACOs* also implies the existence of mechanisms involved in the cell type-specific ethylene response (i.e., root vs RAM shortening). This might be due to connections to spatial-specific signaling circuits (necessarily being different in the differentiated elongated cells and proliferating RAM cells), possibly associated with differential ethylene sensitivity and controlling specific gene sets. This was recently demonstrated in the epidermis, a tissue predominantly controlling ethylene-mediated root and shoot growth (Vaseva et al., 2018). Further, we cannot exclude cell-type specific ethylene distribution reflecting the spatial-specific distribution of ACOs. Considering the gaseous nature of ethylene, this is rather counterintuitive. Nonetheless, oxygen has been implicated as an endogenous diffusible signal in plants in the formation of a hypoxic niche in the shoot apical meristem (SAM) organizing center controlling SAM meristematic activity by regulating *WUSCHEL (WUS)* transcription (Weits et al., 2019). This implies the existence of mechanisms allowing cell type-specific gas distribution in plant tissues as recently demonstrated for tissue-specific regulation of lipid polyester synthesis genes, ensuring a microaerophilic environment in *Lotus* nodules (Venado et al., 2022). That ethylene is insoluble in water may contribute to the possibility of local ethylene action. In parallel, our data imply only limited ability of cytokinins to be transported (either actively or via passive diffusion) within the diverse cell types of the RAM upon the spatially specific upregulation of cytokinin biosynthesis. This is in line with several other reports, suggesting a paracrine mechanism of cytokinins action (Bielach et al., 2012; Bohner and Gatz, 2001), possibly mediated via combined action of cell-type specific cytokinin biosynthesis and degradation (Miyawaki et al., 2004; Waidmann et al., 2019). However, whether there is a cell type-specific ethylene distribution in the *Arabidopsis* root and how it is maintained, remains to be demonstrated.

### Both MSP and canonical ethylene signaling interact in the control of *ACO4*

The mechanisms mediating cytokinin/ethylene crosstalk at the signaling level have been described [for a recent review see (Binder, 2020; Skalak et al., 2021)]. Here we demonstrate the existence of previously uncharacterized signaling mechanism based on a direct interaction between ARR2 and EIN2-C, components of MSP and canonical ethylene signaling, respectively. Our data suggest that both ARR2 and EIN2 are necessary for the ethylene-mediated activation of *ACO4* by cytokinins. ARR2 was found to act downstream of ETR1 in ethylene-dependent signal transduction, possibly mediated via ETR1-dependent ARR2 phosphorylation (Hass et al., 2004). Thus, ethylene might upregulate *ACO4* by activating MSP via ARR2 phosphorylation that recruits the nuclear-localized EIN2-C, a result of the activation of canonical ethylene signaling. Alternatively or additionally, the cytokinin-induced phosphorelay may activate ARR2 by phosphorylation.

Given the positive regulation of *ACO4* by ARR2 and EIN2 *in planta*, our data from the protoplast assay suggesting a negative regulation of *ACO4* in the presence of both EIN2-C and ARR2 seems to be counteracting evidence. However, a similar discrepancy was also previously observed in ethylene- and ARR2-mediated regulation of the ethylene reporter *ERF1:LUC*. In protoplasts, ethylene mediated the downregulation of *ERF1* promoter activity (Hass et al., 2004), while it was upregulated by ethylene in *Arabidopsis* seedlings (Solano et al., 1998). Similarly, in a protoplast assay, the dominantly-active ARR2^D80E^ suppressed *ERF1:LUC* activity even though elevated levels of *ERF1* mRNA were found in *Arabidopsis* seedlings overexpressing *ARR2*^*D80E*^ (Hass et al., 2004). This could be explained by the different developmental context and/or altered responsiveness due to upregulated ethylene production and elevated ethylene signaling in plant protoplasts, associated with upregulation of ethylene-responsive genes [(Hass et al., 2004; Yanagisawa et al., 2003), Jan Lohmann, personal communication]. The abnormal ethylene signaling status of suspension culture-derived protoplasts could also explain their inability to respond to exogenous ACC (this work). The activation of *pACO4* observed in plant protoplasts after overexpression of either *EIN2-C* or *ARR2* alone might be mediated by their interaction with endogenous ARR2 and EIN2-C, respectively. Furthermore, we cannot exclude the effect of the regulatory role of the N-terminal Nramp-like domain of EIN2, which is absent in the EIN2-C construct used in our protoplast assay; it was shown to be necessary for the EIN2-mediated triple response in etiolated seedlings (Alonso et al., 1999). However, in spite of the ambiguities associated with the experimental model used, data from our LUC protoplast assay confirm the ability of EIN2-C and ARR2 to control the activity of the *ACO4* promoter and imply the importance of ARR2 phosphorylation at its conserved Asp 80 in the regulation of *ACO4*.

How the ARR2/EIN2-C complex mediates *ACO4* upregulation is unclear. In canonical ethylene signaling, EIN2-C, which is unable to directly bind DNA, interacts with EIN2 NUCLEAR-ASSOCIATED PROTEIN 1 (ENAP1) leading to the acetylation of histone H3 (H3K14 and H3K23). That induces chromatin to switch to the open state in the ENAP1-binding loci, thus facilitating EIN3-regulated transcription (Wang et al., 2017; Zhang et al., 2016; Zhang et al., 2017). Based on that, one may speculate that ARR2 targets EIN2-C to MSP-regulated loci including *ACO4*, allowing transcriptional activation via EIN2-C-regulated histone acetylation. The mechanism underlying this type of transcriptional activation, nevertheless, remains to be clarified.

### Importance and future outlines

Our findings clearly demonstrate a tight interconnection between cytokinin action and ethylene biosynthesis. Our data imply the existence of a complex network allowing cytokinin control over ethylene biosynthesis at the level of both ACC production and ACC oxidation, the two steps dedicated specifically to ethylene biosynthesis (Depaepe and Van Der Straeten, 2020; Pattyn et al., 2021). Cytokinin-induced *ACSs* and *ACOs* reveal spatial specificity, correlating with the two major roles of ethylene in the control of root growth – regulation of i) cell elongation in the transition/elongation zone and ii) cell division/differentiation in the transition zone/proliferation domain. Our observations also reveal the existence of potential positive feedback regulatory loops, allowing self-potentiation of ACC and ethylene production. Apart from ACS2/6 stabilization by ethylene under stress conditions (Vandenbussche et al., 2012), this type of regulation has been described for ethylene-regulated *ACS* and *ACOs* in the ethylene-induced wilting triggered by pollination in orchids, suggesting that ethylene is not just a switch, but rather is a regulatory factor whose presence is required for a longer period of time [(Dolan, 1997) and references therein]. We found that *ACO3* is a direct target of MSP signaling and described a novel mechanism involving a physical interaction between proteins mediating MSP and canonical ethylene signaling controlling *ACO4* expression. Considering the previously identified integration of both ethylene and cytokinin signals in MSP signaling, this type of regulation represents another level of complexity and control in cytokinin/ethylene crosstalk. Both hormones were shown to control root growth and adaptation by mediating interaction between intrinsic developmental pathways, regulating root development and patterning very early in embryogenesis (Yamoune et al., 2021) and environmental signals (Skalak et al., 2021). This allows the root not only to adapt to immediate conditions including e.g., water availability or soil compaction at the level of root growth and architecture (Chang et al., 2019; Pandey et al., 2021; Park et al., 2018; Saucedo et al., 2012; Szmitkowska et al., 2021; Waidmann and Kleine-Vehn, 2020; Waidmann et al., 2019), but also to anticipate future development and capitalize from past experience via hormone-regulated priming to different stresses (Cortleven et al., 2019; Kosakivska et al., 2022; Skalak et al., 2021; Tiwari et al., 2022). A detailed description of the underlying molecular mechanisms is critical in order to understand the principles activating growth or defense responses in plants and the identification of novel breeding targets. This appears to be highly promising particularly in the era of targeted crop improvement via genome editing approaches.

## Methods

### Plant materials

*Arabidopsis thaliana* ecotype Columbia-0 (Col-0) was used as the Wild Type (WT) and is the background of all the mutants and reporters used in this study. All the T-DNA knockout lines as well as the *pACS*::GUS promoter fusion lines from Tsuchisaka and Theologis (2004 were ordered from NASC. *aco2* “AT1G62380” (N674747), *aco3* ‘‘AT1G12010’’ (N682580), *aco4* ‘‘AT1G05010’’ (N514965), *acs2 “AT1G01480”* (N16564), *acs4* “AT2G22810” (N16566), *acs5-1* “AT5G65800” (N16567), *acs6* “AT4G11280” (N16569), *acs7* “AT4G26200” (N16570), *acs8* “AT4G37770” (N566725), *acs9-1* “AT3G49700” (N16571), *acs5acs9* (N16593), *AmiRacs* (N16651), *arr1-3* “AT3G16857” (N6971), *arr10-1 “AT4G31920”* (N6369), *arr12-1* “AT2G25180” (N6978), *pACS4*:GUS (N31381), *pACS5*:GUS (N31382), *pACS6*:GUS (N31383), *pACS8*:GUS (N31385), *pACS9*:GUS (N31386) and *pACS11*:GUS (N31387), *acs2-1acs4-1acs5-2acs6-1acs7-1acs9-1amiRacs8acs11* (*acs8x*; Tsuchisaka et al., 2009), *eto1-1* (Woeste et al., 1999) was procured from ABRC. The double mutant *aco2aco3* and *aco2aco4* were generated by crossing the corresponding single mutants.

The ectopic cytokinin overproducing lines were prepared using the GAL4>>UAS-based two-component activator-reporter system, J2601 and J2351 activators were respectively crossed to the reporter UAS::IPT for ectopic IPT overexpression, or to the Col-0 as controls. F1 generation of the crosses was used for analysis, as they were sterile. J2601, J2351 and UAS::IPT (Laplaze, et al. 2007) lines were kindly provided by Prof. Eva Benkova (Bielach et al. 2012).

All the fluorescent reporter lines produced in this study were generated by the flower dip method as described by Clough and Bent (1998) in the Col-0 background, and single copy homozygous T3 lines were selected and used for analysis. The *arrB-pACO3*::ACO3:GFP and *arrB-pACO4*::ACO4:GFP and *ein2-1-pACO4*::ACO4:GFP lines were generated by crossing the single *arrB* (*arr1-3, arr2-5, arr10-1, arr12-1)* mutants or *ein2-1* (Zdarska et al., 2019) respectively with the generated reporter lines *pACO3*::ACO3:GFP and *pACO4*::ACO4:GFP.

### Growth conditions

Seeds were surface sterilized and sown on half-strength Murashige–Skoog (½MS) medium (Duchefa Biochemie) with 1% (W/V) sucrose and 1% (W/V) plant agar, then stratified in the dark for 2 days at 4°C. seedlings were grown vertically under long-day-conditions (16 h light/8 h dark) at 22°C for the duration of the treatment.

### Cloning

Unless otherwise specified, all the cloning was done using the Gateway™ (Invitrogen) system following the manufacturer’s instructions. Fragments were isolated by PCR amplification using Phusion® High-Fidelity DNA Polymerase (NEB) from Col-0 genomic DNA/cDNA. For every step of the cloning, all cloned sequences were verified by colony PCR, plasmid digestion and sequencing. The primers used are described in supplemental material table 1.

### Fluorescent reporters

Entry clones of the native promoters (∼2.5Kb upstream ATG) and/or promoter-coding sequence (as one fragment without stop codon), of *ACS2, ACS7* and/or *ACO1,3-5*, were cloned either in pFAST-G04 for transcription and/or pFAST-R07 for translation fusions (Shimada et al., 2010) respectively. The *pACO2*::ACO2:GFP clone was prepared by replacing the 35S promoter, in the p2GWF7.0 vector (Karimi et al., 2007), with the native *ACO2* promoter using Gibson Assembly® (NEB) following the manufacturer’s instructions, the coding sequence was cloned afterwards by LR reaction. The generated clones are: pACO3/4::GFP:GUS, pACO1-5::ACO1-5:GFP and pACS2,7::ACS2,7:GFP.

### Y1H assay

DNA-bait and prey clones were generated as described by Reece-Hoyes and Walhout (2018 a, b). Overlapping bait promoter fragments of *ACO3* and *ACO4* promoters (Fig. 4) were each cloned into pDONR-P4P1r, then respectively into pMW#2 and pMW#3 to generate (pACO3/4::HIS and pACO3/4::LacZ). For the cDNA prey clones, the entry clones pENTR_ΔDDK-ARRB and pENTR_EIN2-C, generated by cloning into pDONR221 the truncated type-B ARRs missing the response regulator domains (ΔDDK-ARR1,2,10,12 see Suppl. Fig. 4) and EIN2-Cend, were cloned into pDEST22 for ΔDDK-ARRB (AD-ΔDDK-ARRB) and pGADT7 for EIN2-C (AD-EIN2-C).

For the FLIM-FRET assay, pENTR-ΔDDK-ARR2 and pENTR-EIN2-C (from Y1H) were respectively cloned into the pB7WGR2 and pH7WGF2 destination vectors (Karimi, et al. 2007) for overexpression and N-terminus fusions to both GFP and RFP (35S::GFP:ARR2, 35S::RFP:ARR2, 35S::GFP:EIN2-C, and 35S::RFP:EIN2-C). As a positive control the binary vector 35S::GFP:NLS:RFP was constructed by fusing the NLS:RFP to the GFP in pH7WGF2 destination vector (Karimi et al., 2007) via LR reaction, RFP isolated from pB7WGR2 to generate pENTR-NLS:RFP by BP; the NLS sequence was added into the forward nls::RFP-attB1-F primer.

### Luciferase assays

The *pACO4::LUC* reporter lines were prepared by cloning a 2547 bp (−2232 to +315) long fragment promoter region into the *pBT8-LUCm*^*3*^ (Wallmeroth et al., 2019) via EcoRI and NcoI restriction sites. The effector cDNA of the C-terminal fragment of EIN2 (F646 to G1294) was cloned into the *pHBTL-3xHA-GW* (Ehlert et al., 2006). The effector cDNA *ARR2, ARR2*^*D80E*^ and *ARR2*^*D80N*^ in the *pHBTL-3xHA-GW* vector (Wallmeroth et al., 2019) were used. *ARR2*^*D80A*^ was obtained from the *ARR2* cDNA by site-directed mutagenesis (QuikChange Lightning Site-Directed Mutagenesis kit – Agilent).

## Supporting information

Yamoune et al., supplemental information

## Author contributions

A.Y., M.Z and J.H. conceived the research; A.Y., M.Z., T.D., A.K., E.S., K.B., V.M-R., J.S., P.T., L.B., B.P., and A.C. performed the research; A.Y., M.Z., T.D., K.B., V. M-R., J.S., L.B., I.K., M.P., O.N., E.M., K.H., D. VDS and J.H. analyzed the data; A.Y., M.Z., E.M., K.H., D. VDS. and J.H. wrote the paper.

## Acknowledgements

We are very grateful to prof. Li-Jia Qu for providing us with seeds of ethylene-free lines (Li et al., 2022). We acknowledge the core facility CELLIM of CEITEC supported by the MEYS CR (LM2018129 Czech-BioImaging). Core Facility Plants Sciences of CEITEC MU is gratefully acknowledged for obtaining scientific data presented in this paper. This article is subject to HHMI’s Open Access to Publications policy. HHMI lab heads have previously granted a nonexclusive CC BY 4.0 license to the public and a sublicensable license to HHMI in their research articles. Pursuant to those licenses, the author-accepted manuscript of this article can be made freely available under a CC BY 4.0 license immediately upon publication.

