## Supplementary material for "Cytokinins regulate spatially-specific ethylene production to control root growth in *Arabidopsis*": Yamoune et al., supplemental information

**Supplemental Figures**

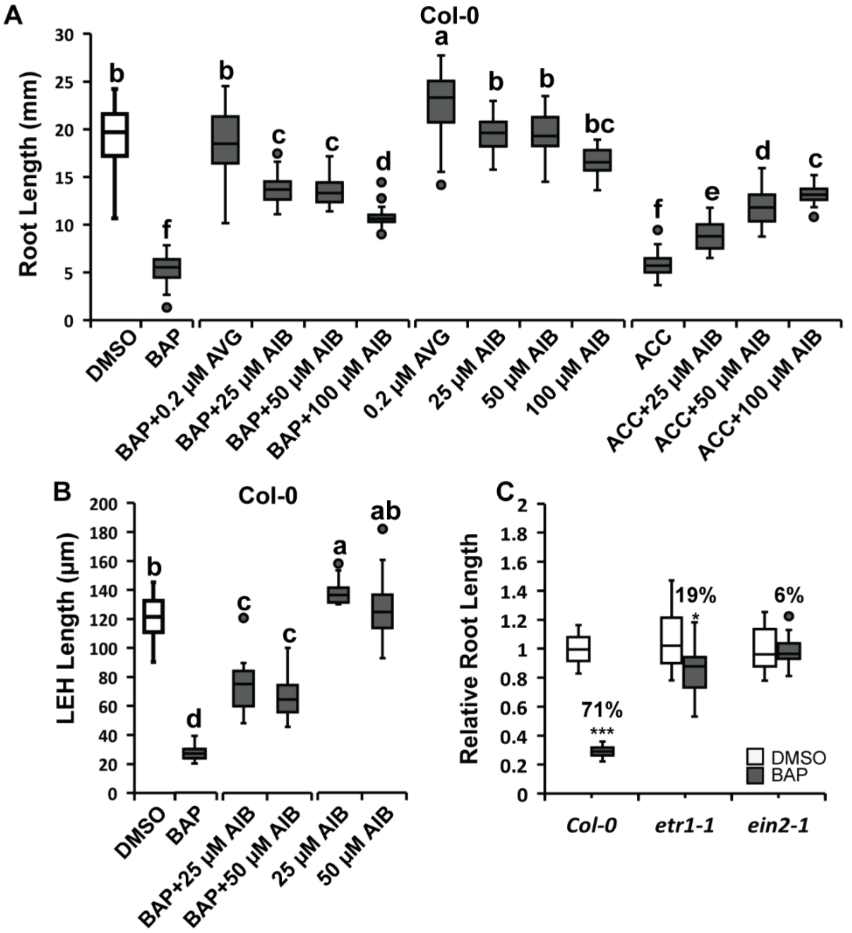


**Supplemental Figure 1. Cytokinin-induced root shortening is mediated by ACC/ethylene biosynthesis and ethylene signaling**

(**A**) Root length of six-day-old *WT Col-0* seedlings grown on ½MS +/- 0.1 µM BAP combined with AVG or AIB and their respective controls or on +/- 1 µM ACC ½MS with or without AIB. (**B**) Length of the first Epidermal cell with a visible root Hair bulge (LEH) of six-day-old *WT Col-0* seedlings grown on ½MS + 0.1 µM BAP with or without AIB. Boxplots represent data from the three independent replicates. The letters represent significance classes determined by one-way ANOVA followed by Tukey's post-hoc HSD test. (**C**) Relative root length of six-day-old *WT Col-0*, *etr1-1* and *ein2-1* seedlings grown on 0.1 µM BAP (control is 0.01% DMSO). Boxplots represent root length normalized to the respective mock-treated control (DMSO), * or *** denote the Student's t-test significance at p <0.05 or p <0.001 respectively.

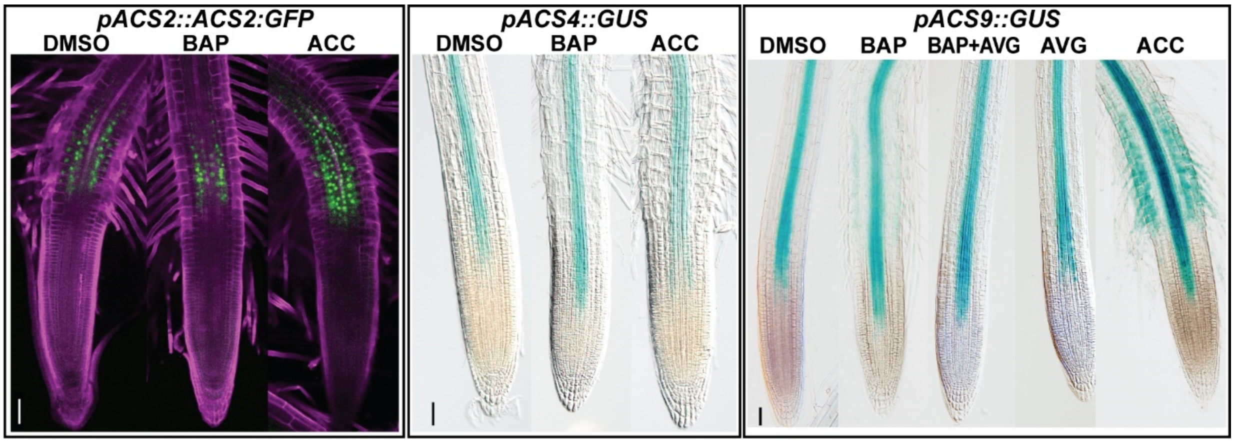


**Supplemental Figure 2. Cytokinin non-responsive *ACS* genes**

Six-day-old *pACS2*::ACS2:GFP, *pACS4*::GUS and *pACS9*::GUS reporter seedlings treated for 24h with 5 µM BAP, 5 µM BAP + 1 µM AVG, 1 µM AVG, or 5 µM ACC (control is 0.01% DMSO). The scale bars represent 50 µm.

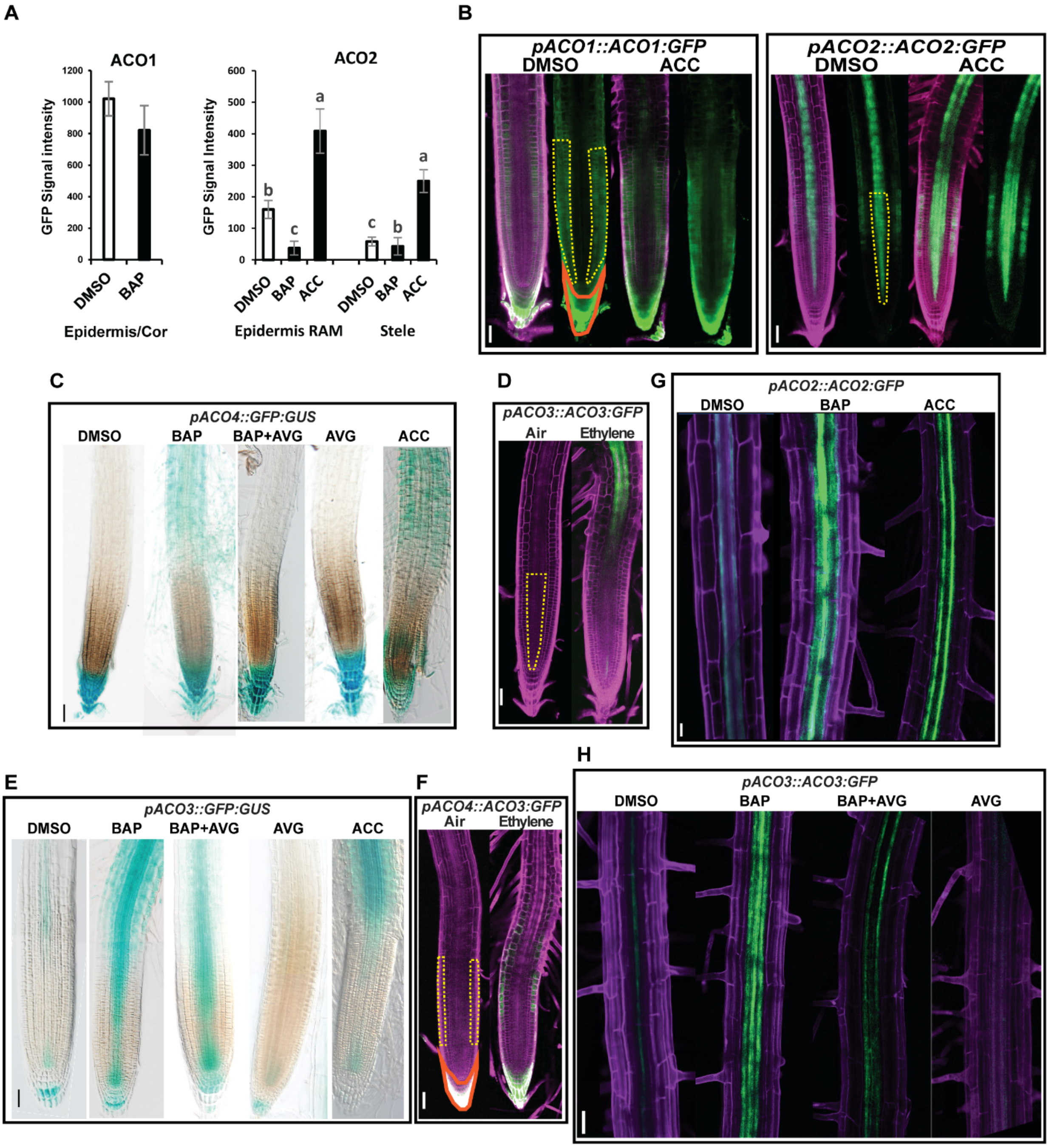


**Supplemental Figure 3. Promoter activity and protein quantification of *ACC oxidase* genes.**

(**A**) Relative GFP signal quantification and (**B**) ACC-mediated induction of *ACO1* and *ACO2* respectively seen in the root tips of six-day-old seedlings of *pACO1::ACO1:GFP* (left) and *pACO2::ACO2:GFP* (right) treated for 24h with 5 µM BAP or 5 µM ACC; control is 0.01% DMSO. Bars represent means ± SD and the letters significance classes (one-way-ANOVA followed by Tukey's post-hoc HDS test). (**C**) Hormonal control of *ACO3* and (**E**) *ACO4,* visualized by GUS staining of six-day-old *pACO3::GFP:GUS* and *pACO4::GFP:GUS* translation fusion lines respectively, treated for 24h with 5 µM BAP, 5 µM BAP+0.2 µM AVG, 0. 2 µM AVG or 5 µM ACC; control is 0.01% DMSO. (**D**) Six-day-old *pACO3::ACO3:GFP* and (**F**) *pACO4*::ACO4:GFP seedlings treated for 24h with 10 ppm of ethylene with air as control. (**G**) Root maturation zone of six-day-old seedlings of *pACO2*::ACO2:GFP and (**H**) *pACO3*::ACO3:GFP treated for 24h with the indicated hormones (5 µM BAP, 5 µM BAP+0.2 µM AVG, 0.2 µM AVG or 5 µM ACC; control is

0.01% DMSO). The dotted lines in B, D, and F mark the area in which the GFP signal was quantified. The scale bars (B-F, H) represent 50 µm, and 20 µm in (G).

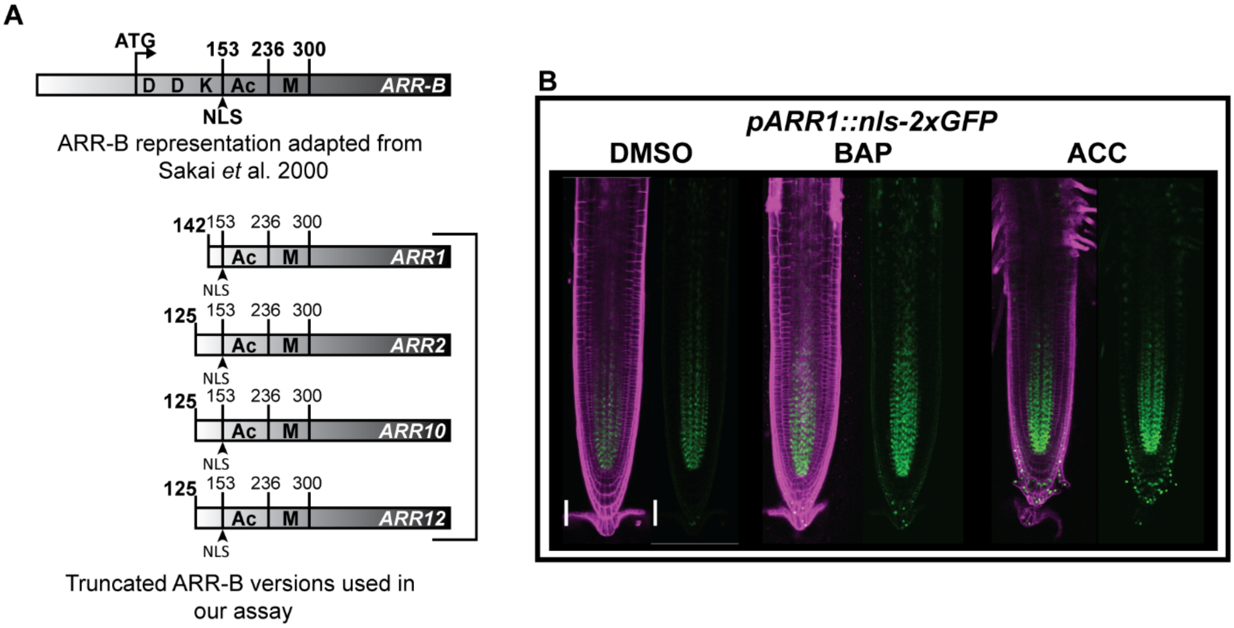


**Supplemental Figure 4. Type-B ARRs are under hormonal control and control promoter activity of *ACO3* and *ACO4.***

**(A)** Schematic representation of the truncated versions of the type-B ARRs [based on (Sakai et al., 2000b)] cloned for the Y1H assay. **(B)** Both cytokinins and ACC upregulate *ARR1* in the stele. The scale bars (B) represent 50 µm.

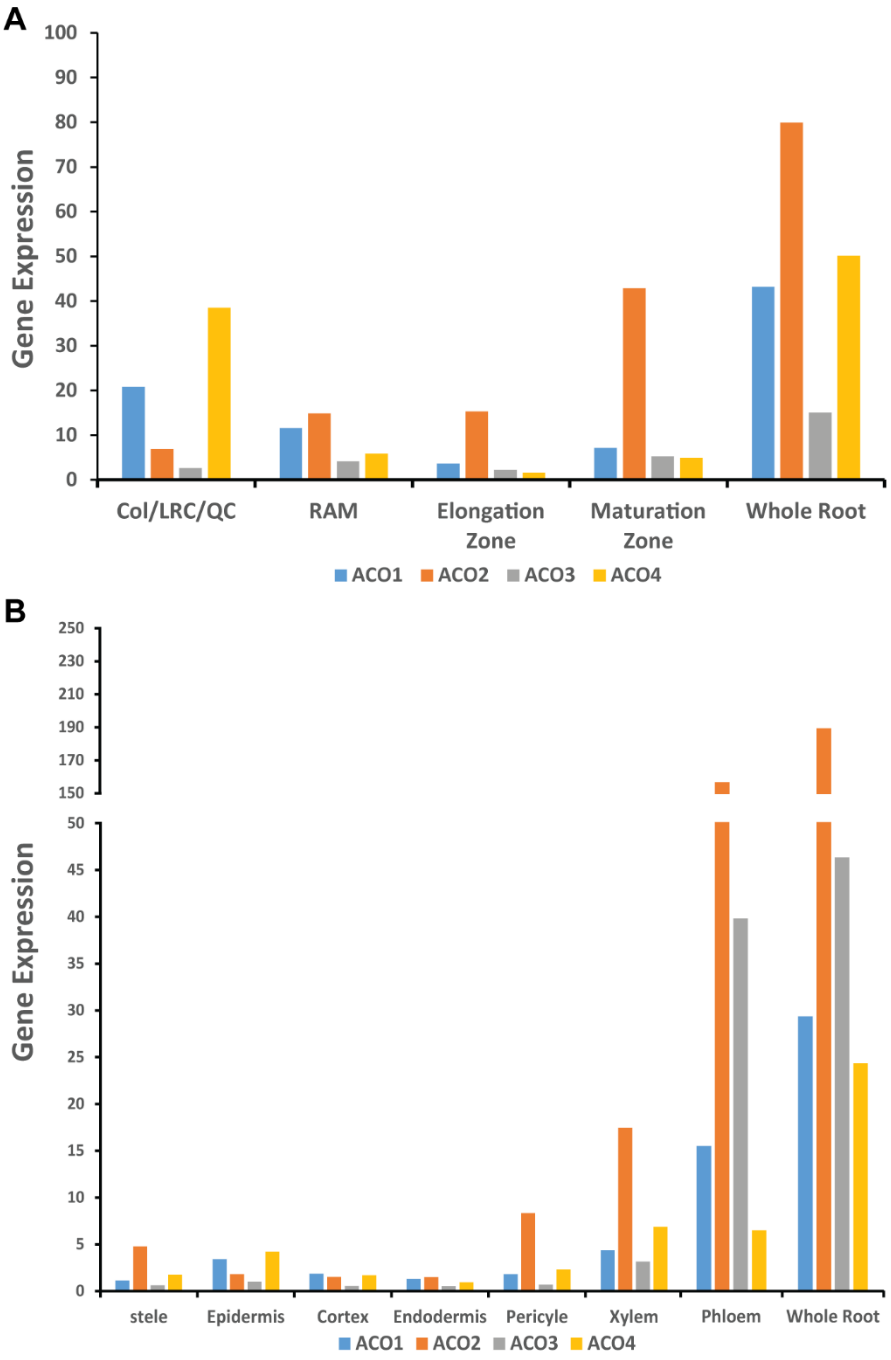


**Supplemental Figure 5. *ACO* gene family expression in the *Arabidopsis thaliana* root.**

Expression of different ACO genes (*ACO1*-4) in the root of *Arabidopsis thaliana* in the longitudinal **(A)** and the radial **(B)** axes. Data based on Brady et al. (2007) and were extracted from http://bar.utoronto.ca/eplant/. While *ACO2* is rather poorly expressed in the columella/LRC/QC, it is the *ACO* with highest expression levels in the rest of the root.

### Supplemental Methods

#### Hormonal treatments

Depending on the type of analysis we used two types of assays. Direct treatments were used for the root and LEH length, where seeds were sown directly and grown continuously on media containing BAP (B3408, Sigma-Aldrich), ACC (A3903, Sigma-Aldrich), AIB (850993, Sigma-Aldrich), or AVG (32999, Sigma-Aldrich). For the RAM and reporter lines analysis, seeds sown and grown on ½MS, were transferred to liquid ½MS supplemented with the different hormonal treatments.

#### GUS staining

Six-day-old pACS::GUS and pACO3-4::GFP:GUS seedlings were stained for GUS expression as described in Malamy and Benfey (1997). Differential Interference Contrast microscopy (Olympus BX61) was used for imaging. 1 mM of Fe salts (K3, K4) were used. pACS::GUS seedlings were stained for 15 min and pACO::GFP:GUS ones for 30 min .

#### Root and RAM visualization, measurements and reporter image analysis

Roots were stained with propidium iodide (PI; P4864, Sigma-Aldrich; 50µg per ml of H2O for 7min) and imaged using the inverted confocal microscopy system Zeiss LSM 800 or LSM 880 to visualize root cells. RAM size was determined according to Dello Ioio *et* al. (2007), as the number of the cortex cells counted from the quiescent center to the first elongated cell. The length of the cortical cells was first measured using Cell-O-Tape macro (French et al., 2012) in ImageJ/Fiji software (Schindelin et al., 2012) (http://rsb.info.nih.gov/ij/) and the first elongated cell was designated by performing a point change test of the cells length using "The Multiple Structural Change algorithm" tool available at http://www.ibiologia.com.mx/MSC_analysis (Pacheco-Escobedo et al., 2016). The first epidermal cell with visible root hair bulge (LEH) was determined as described in Le *et* al. (2001). Both LEH and root lengths were measured using ImageJ/Fiji software. For root elongation, plates were scanned with the same ruler to set the scale.

#### ACC Measurement

50 root tips from each sample were cut and frozen in cold water and assessed for ACC levels. ACC quantification was performed using a modified version of the method previously published by (Salazar et al., 2012). Briefly, roots of *Arabidopsis thaliana* were extracted into 1ml of $H_2O$:methanol (1:1) extraction solution and $d_4$-ACC was added as an internal standard. The samples were homogenized using a bead mill (MixerMill, Retsch GmbH, Haan, Germany) and then centrifuged for 15 minutes at 18 000 rpm at 4°C. 400 µL of supernatant was evaporated

to dryness and derivatized with the AccQ-Tag Ultra kit (Waters, Milford, MA, USA). For ACC quantification, the samples were analyzed by liquid chromatography–tandem mass spectrometry (LC-MS/MS) in multi reaction monitoring (MRM) mode employing the LC-MS/MS system 1260 Infinity II LC System coupled to a 6495 Triple Quad LC/MS System with a Jet Stream and Dual Ion Funnel technologies (Agilent Technologies, Santa Clara, CA, USA).

**Quantification of Reporter Gene Expression**

Quantification of the pACO1-4::ACO1-4:GFP signal was performed in Fiji as described in Zdarska & Cuyacot (2019), the region of interest for each gene was defined as in supplemental figure 3.

***In vivo* Ethylene Measurement**

Six-day-old Col-0, *aco2*, *aco3*, *aco4*, *aco2aco3,* and *aco2aco4* seedlings were grown vertically on a mesh on ½MS (1% (w/v) sucrose solidified with 0.8% (w/v) agar. The roots were then cut and wound ethylene was allowed to dissipate for 4h on the plate (to preserve humidity). Then 30 roots from each of the tested genotypes were subsequently transferred to 10 mL chromatography vials (Chromacol, VWR) with 4 mL liquid ½MS + 5 µM BAP or 0.01% DMSO as a control. The vials were hermetically sealed with rubber stops and snap-caps (Chromacol, VWR). Roots were incubated for 48h with gentle shaking to allow ethylene accumulation in the headspace. Ethylene emanation was measured using laser-based photoacoustic spectroscopy (ETD-300, Sensor Sense, The Netherlands) according to Van de Poel & Van Der Straeten (2017). Three biological replicates were measured; five vials containing 30 seedlings each, were used for each biological replicate and each treatment/genotype combination. 4 blank vials, containing only medium were used as controls for the measurements. Data were normalized to the blank control and expressed per hour per mg fresh weight.

**Yeast transformation, mating and Y1H screening**

The DNA-bait clones were integrated into YM4271 yeast strain genome and the AD-TF transformed into Y187α as described by Reece-Hoyes and Walhout (2012). AD-ARR2 and AD-EIN2-C were co-transformed as well. To screen AD-TF/bait-HIS3 activation, mating was allowed as described in Castrillo et al (2001). The diploid-selective-media used were: CSM-His-Ura-Trp for AD::ARRs/bait-HIS3; CSM-His-Ura-Leu for EIN2-C/bait-HIS3, and CSM-His-Ura-Trp-Leu for AD::ARR2+AD::EIN2-C/bait-HIS3. The screening plates contained the diploid-selection-media ± an ascending concentration of 3-AT (0 mM, 20 mM, 40 mM, 60 mM, and 80 mM). The YM4271 and Y172α yeast strains were kindly provided by Helene Robert Boisivon, PhD.

**FLIM-FRET**

Plasmid vectors (35S::GFP:ARR2, 35S::RFP:ARR2, 35S::GFP:EIN2-C, and 35S::RFP:EIN2-C as well as the positive control 35S::GFP-RFP) were transiently expressed in *Nicotiana tabacum* (SR1 Petit Havana) epidermal leaf cells using the infiltration procedure described in (Voinnet et al., 2000). Gene silencing in *Nicotiana tabacum* was suppressed by co-infiltrating the p19 protein from tomato bushy stunt virus cloned into pBIN61 (Voinnet et al., 2000). The Zeiss LSM 780 Axio-Observer laser scanning confocal imaging microscope equipped with external In Tune laser (488-640 nm, < 3nm width, pulsed at 40 MHz, 1.5 mW) C-Apochromat 63 x water objective, NA 1.2 and the HPM-100-40 Hybrid Detector from Becker and Hickl GmbH was used for FLIM-FRET data acquisition. FLIM analysis was performed using a Simple-Tau 150N (Compact TCSPC system based on SPC-150N) with DCC-100 detector controller for photon counting. For GFP and RFP excitation, we used the Tune laser at 490 nm wavelength and a DPSS-laser at 561 nm, respectively. Zen 2.3 light version from Zeiss was used for processing confocal images. SPCM 64 version 9.8 was used to acquire FLIM data and SPCImage version 7.3 from Becker and Hickl GmbH for data analysis. For each analysis, the nuclear area was selected as region of interest containing signals for fluorescent lifetime calculation. A multiexponential decay model was used for fitting. Lifetime components with very low values below 500 ns were considered as background and avoided for average lifetime calculations.

**Luciferase assays**

96-well transfection assays for assessing reporter gene expression were essentially performed as in Wallmeroth (2019). In each transfection 30µL of cell-culture protoplasts ($3.5x10^6$ protoplasts $mL^{-1}$ ) for a total of $1.05 \times 10^5$ protoplasts were transfected with a plasmid DNA mixture containing 3 µg *ACO4p::fLUC* reporter-gene plasmid, 2µg pCF203 encoding *35S::eGFP* for normalization, 2 µg of one of the *ARR2* cDNA variants or in combination with 2 µg EIN2c in *pHBTL-3xHA* or substituted with pUC19 as stuffer plasmid to a total of 9 µg in a total of 20 µL water. Cells were transfected with 30 µL PEG1500, the reaction stopped with 30 µL MM and incubated with the addition of 350 µL K3 for a total of 460 µL. A total of four biological replicates per combination were pipetted separately per experimental run. After an overnight incubation, the promoter activity was measured *in vivo* for three hours in 150µL transfected protoplasts from two technical replicates using a Mithras LB 940 Multimode Microplate Reader (Berthold). GFP emission was measured in 100 µL cells in a black microtiter plate with a TECAN Safire (ex488±7.5nm, em525±12nm). Data extraction and statistics were performed in JMP16 SAS. The luciferase emission was normalized to the GFP emission per individual transfection, the peak LUC/GFP emission was extracted, and the average from the technical splits taken for statistical evaluation. Total LUC activity, either summed or calculated as area-under-the-curve, yielded equivalent results as the maximum value (not shown).

**Gene expression analyses**

The root tips from 6-DAG seedlings treated for 6 hours (5µM BAP with 0.01% DMSO as a control) were cut off with a scalpel and immediately frozen in liquid nitrogen. Total RNA from the collected tissue was isolated using the RNAqueous Small Scale Phenol-Free Total RNA Isolation Kit (Ambion) according to the manufacturer's instructions. cDNA was prepared using RTP3 primer and Superscript III (Invitrogen) according to the manufacturer's instructions, and RT–qPCR was performed using the FastStart SYBR Green Master Kit (Roche) according to the manufacturer's instructions on a Rotor-Gene 6000 (CORBETT RESEARCH) instrument. RTP3 primer: 5'-CGTTCGACGGTACCTACGTTTTTTTTTTTTTTTTT-3'. Four independent replicas were processed. For relative quantification of *ACO1-5* transcripts, we used the primers in table 2 (*ACO3* and *ACO5* as described in (Schellingen et al., 2014) and *ACO1-2* and *ACO4* as described in (Lee et al., 2017). For individual *ACOs* pairs of primers sequences are in the Supplemental material table 1. Average relative quantities were normalized to internal controls UBIQUITIN-CONJUGATING ENZYME 10 (UBC10) and to mock-treated controls that correspond to value 1 (qbase+, Biogazelle). For statistical analysis, a Mann–Whitney test was performed (qbase+, Biogazelle).

**Statistics**

All charts (unless otherwise mentioned) were drawn using Microsoft Excel 365®. The statistical analysis, pairwise comparisons, post hoc tests were performed in RStudio. For RAMs, LEH lengths, root lengths, ethylene measurements data were treated as follows: the outliers were detected using the boxplot method, where the data were divided into Genotype and Treatment condition sub-sets. For each subset values above Q3[1] + 3xIQR[2] or below Q1 - 3xIQR were considered extreme points and were removed before further analysis. The effect of different treatments and genotypes on ethylene production was assessed by fitting a linear mixed effect model via restricted maximum likelihood. The experiment was carried out with 2 or 3 biological replicates which was introduced as a random effect in the model to account for intra-cluster correlation between the responses from one replicate. The model was defined considering the interaction between genotype and treatments. Pairwise comparison was carried out using Kenward-Roger degrees-of-freedom calculation and Tukey's adjustment. For ethylene measurements, the pairwise comparison of genotype was calculated independently for each treatment group, as the result of the mixed model stated the differences between treatments clearly ($p<0.001$). For all the other measured characteristics pairwise comparisons were computed among genotypes and treatments together. The *lmer* and *emmeans*[3] packages of software R were used to calculate the results (Bates et al., 2015; R_Core_Team, 2021).

[1] 3rd quartile
[2] inter quartile range
[3] Lenth, Russell V., Paul Buerkner, Maxime Herve, Maarten Jung, Jonathon Love, Fernando Miguez, Hannes Riebl, a Henrik Singmann. "emmeans: Estimated Marginal Means, aka Least-Squares Means", 08. september 2022. https://CRAN.R-project.org/package=emmeans

**Supplemental Table 1:** List of primers used.

| Gene | PRIMER | SEQUENCE | AIM |
|---|---|---|---|
| **ACO1** | pACO1-attB1-F<br>ACO1-attB2-R | GGGGACAAGTTTGTACAAAAAAGCAGGCTTAccggtggttggagaacgaga<br>GGGGACCACTTTGTACAAGAAAGCTGGGTTCggctgaatccgcatttccca | BP: pDONR221 then pFAST-R07 (Protein) |
| | ACO1-qRT-F<br>ACO1-qRT-R | TTGAGTGAAGGCAAAACCTCAGATG<br>GCTGAGTTCCTCTGAAATGTTTGGG | RT-qPCR (Lee et al., 2017) |
| **ACO2** | pACO2-F<br>pACO2-R | CAAGCTAAGCTTGAGCTCTATCAATTATTTCTCGTGGTTTTTTG<br>TTTTTGTACAAACTTGTGATATCACTAGTCTTTCTTTCTCTCTCTCTTCTTTGA | Gibson assembly to replace 35S in p2GWF7.0 |
| | ACO2- attB1-F<br>ACO2- attB2r-R | GGGGACAAGTTTGTACAAAAAAGCAGGCTTAatggagaagaacatgaagtttccag<br>GGGGACCACTTTGTACAAGAAAGCTGGGTTCgaaagtctctacggctgctgtag | BP: pZeo then modified p2GWF7.0 |
| | ACO2-LP<br>ACO2-RP | GACAATCACAGCTGAGGAAGC<br>TTAAACCGGAAGAACGACATG | genotyping / sequencing |
| | ACO2-qRT-F<br>ACO2-qRT-R | GCACCGTGTGGTGACTCAACA<br>AAGTCTCTACGGCTGCTGTAGGAT | RT-qPCR (Lee et al., 2017) |
| **ACO3** | pACO3-attB1-F<br>pACO3-attB2-R<br>ACO3- attB2-R | GGGGACAAGTTTGTACAAAAAAGCAGGCTTAgaggtctccgcattggggttg<br>GGGGACCACTTTGTACAAGAAAGCTGGGTTCtctctctctctctcttaactagctact<br>GGGGGACCACTTTGTACAAGAAAGCTGGGTTCgaatgtctcaaccacagccacc | BP: pDONR221 then pFASTG04 (Promoter) pFAST-R07 (Protein) |
| | ACO3-LP<br>ACO3-RP | ATCCCATCTCAAAAGCAGGAG<br>CTTGAAACAGCAAAATGAGGC | genotyping / sequencing |
| | ACO3-qRT-F<br>ACO3-qRT-R | CAAGCATTCCATTGTCATCAACCTTG<br>TTTCTGGGTCATCACACGGTG | RT-qPCR (Schellingen, et al. 2014) |
| | pACO3-1-attB4-F<br>pACO3-1-attB1r-R | GGGGACAACTTTGTATAGAAAAGTTGCTggaatttgtctcctcatcctgcta<br>GGGGACTGCTTTTTTGTACAAACTTGTctctctctctctcttaactagctact | BP: pDNOR221 P4-P1r, For Y1H-BAIT: pPMW#2, pMW#3 |
| | pACO3-2-attB4-F<br>pACO3-2-attB1r-R | GGGGACAACTTTGTATAGAAAAGTTGCTttgtaattatattagctggccaagg<br>GGGGACTGCTTTTTTGTACAAACTTGTtatttgaatcactatgaatagggaatgac | |
| | pACO3-3-attB4-F<br>pACO3-3-attB1r-R | GGGGACAACTTTGTATAGAAAAGTTGCTcgttaccattgaaagtaagtattttgttca<br>GGGGACTGCTTTTTTGTACAAACTTGTaactttgttaattttgggaaggaag | |
| | EcoRI-pACO3 fw<br>NcoI-pACO3 rev | TCTATTATC GAATTC GAGGTCTCCGCATTGGGGTTG<br>TCTATTATC CCATGG CTCTCTCTCTCTCTCTTAACTAGCTACT | LUC fusion |
| **ACO4** | pACO4-attB1-F | GGGGACAAGTTTGTACAAAAAAGCAGGCTTAtccgcggattctatcttcgtacttgcat | BP: pDONR221 then |

| | | | |
|---|---|---|---|
| | pACO4-attB2-R<br>ACO4- attB2-R | GGGGACCACTTTGTACAAGAAAGCTGGGTTCtctctctctttttttttaaatgggtttcttg<br>GGGGACCACTTTGTACAAGAAAGCTGGGTTCcgcagtggccaatggtcc | pFASTG04 (Promoter)<br>pFAST-R07 (Protein) |
| | pACO4-Seq1-F<br>ACO4-Seq2-R | GACTTCTCAAGTTGTTGTTTTGTA<br>AAAAGGTTACCTGTAATCGTCG | sequencing |
| | ACO4-LP<br>ACO4-RP | GTCCATATGCATTTGGACTGG<br>GGAGCTACTGGATCTGCTGTG | genotyping / sequencing |
| | ACO4-qRT-F<br>ACO4-qRT-R | GAGTGCTATCTCAGACAGACGGAG<br>CTTGGTTCCTTGGCCTGAAACTTG | RT-qPCR (Lee et al., 2017) |
| | pACO4-1-attB4-F | GGGGACAACTTTGTATAGAAAAGTTGCTggaagaaaacgggtcaaacaatg | BP: pDNOR221 P4-P1r,<br>For Y1H-BAIT: pPMW#2, pMW#3 |
| | pACO4-1-attB1r-R | GGGGACTGCTTTTTTGTACAAACTTGTtctctctctctttttttttaaatgggtttcttg | |
| | pACO4-2-attB4-F | GGGGACAACTTTGTATAGAAAAGTTGCTataacaaagtatgaatgttgatcaaagaca | |
| | pACO4-2-attB1r-R | GGGGACTGCTTTTTTGTACAAACTTGTgggtcggaaaaaatataaaaaatttatg | |
| | pACO4-3-attB4-F | GGGGACAACTTTGTATAGAAAAGTTGCTaggaggtccactagtaggtcaagtt | |
| | pACO4-3-attB1r-R | GGGGACTGCTTTTTTGTACAAACTTGTcattcattgccttacttcctttcta | |
| | EcoRI-pACO4 fw | TCTATTATC GAATTC TCCGCGGATTCTATCTTCGTACTTGCAT | LUC fusion |
| | NcoI-pACO4 rev | TCTATTATC CCATGG CTCTCTCTCTTTTTTTTTAAATGGGTTTCTTG | |
| **ACO5** | pACO5-attB1-F<br>pACO5-attB2-R<br>ACO5-attB2-R | GGGGACAAGTTTGTACAAAAAAGCAGGCTTA cttgactagttggtatttacgctga<br>GGGGACCACTTTGTACAAGAAAGCTGGGTTCttcagatccgcaaagagagaga<br>GGGGACCACTTTGTACAAGAAAGCTGGGTTCgagagactttacagctagaaaacga | BP: pDONR221 then pFASTG04 (Promoter)<br>pFAST-R07 (Protein) |
| | ACO5-GK-RP<br>ACO5-GK-LP | CCTTTAGGCAAACCCAAATTC<br>TGTAAGGGATTTCTGTTTCATCC | genotyping |
| | ACO5-qRT-F<br>ACO5-qRT-R | TGTTCAGCCTCTACCTAATGCCA<br>CCTGTGCCACGCACTCTTGTA | RT-qPCR (Schellingen, et al. 2014) |
| **ARR1** | arr1-3-F<br>arr1-3-R | CTTCAAGCACTAGCCGTCACAGGTCAGTT<br>AATGTTATCGATGGAGTATGCGTCAAAGT | genotyping |
| | ARR1-142aa-attB1-F | GGGGACAAGTTTGTACAAAAAAGCAGGCTTAatggaggcacttaagaacatatggc | BP: pDONR221 for Prey-pDEST22 (Y1H) |
| | ARR1-Stop-attB2-R | GGGGACCACTTTGTACAAGAAAGCTGGGTTTCaaaccggaatgttatcgatgg | |
| | ARR1 Prom F-Mlu1<br>ARR1 Prom R-EcoR1/EcoRV | cccacgcgtgagtacagctgtgaaattgatggattactacc<br>cccGAATTCGATATCacctctctctatgtagctcgaacccaag | Cloning of *pARR1::nls-2xGFP* |

| | | | |
|---|---|---|---|
| | ARR1 3'UTR R-EcoR1/Stu1 | cccgaattcaggcctttggataaaaaaaccgataaaccggagggact | |
| | ARR1 3'UTR R-Nhe1 | cccgctagcTGGCAAGTCGTCTGCAACATCACTCAACCAAC | |
| **ARR2** | ARR2-5F<br>ARR2-5R | CCTTCTCTGATCGTTCGTTTTCTG<br>ATCAACGACAAGAACTCGAAGATTC | genotyping |
| | ARR2-125aa-attB1-F<br>ARR2-stop-attB2-R | GGGGACAAGTTTGTACAAAAAAGCAGGCTTAatggattacctcatcaaaccggtac<br>GGGGACCACTTTGTACAAGAAAGCTGGGTTTCagacctggatattatcgatggagta | BP: pDONR221 for Prey-Y1h: pDEST22 & FLIM-FRET |
| | ARR2 D80A fw<br>ARR2 D80A rev | ggttttgatattgtcattagtgCtgttcatatgcctgacatgg<br>ccatgtcaggcatatgaacaGcactaatgacaatatcaaaacc | site-directed mutagenesis at D80A site. |
| **ARR10** | arr10-1-F<br>arr10-1-R | GCCACCTTCAGGTGAGAGTTAGACTATGAT<br>AGCTGACAAAGAAAAGGGAAAATGGAGTTT | genotyping |
| | ARR10-122aa-attB1-F<br>ARR10-stop-attB2-R | GGGGACAAGTTTGTACAAAAAAGCAGGCTTAatggaggagcttaagaacatatggc<br>GGGGACCACTTTGTACAAGAAAGCTGGGTTTCaagctgacaaagaaaagggaaaa | BP: pDONR221 for Prey-pDEST22 (Y1H) |
| **ARR12** | arr12-1-LP<br>arr12-1-RP | CGGTACAATATGCGGATTTTGATTCGGTAT<br>TAATAGCTTGCTGATTAGCCACACCACTGA | genotyping |
| | ARR12-122aa-attB1-F<br>ARR12-stop-attB2-R | GGGGACAAGTTTGTACAAAAAAGCAGGCTTAatggaggagttgaagaacatatggca<br>GGGGACCACTTTGTACAAGAAAGCTGGGTTTCatatgcatgttctgagtgaactaaac | BP: pDONR221 for Prey-pDEST22 (Y1H) |
| **EIN2-C** | EIN2-C-459aa-attB1-F<br>EIN2-C-attB2-R | GGGGACAAGTTTGTACAAAAAAGCAGGCTTAatgacgccgctgaaatctgcga<br>GGGGGACCACTTTGTACAAGAAAGCTGGGTTTCaacccaatgatccgtacgcag | BP: pDONR221 for prey-Y1H:pGAT7 and FLIM-FRET |
| **ACS2** | pACS2-attB1-F<br>ACS2-attB2-R | GGGGACAAGTTTGTACAAAAAAGCAGGCTTAgacatgatcactgtgaagtcgtgc<br>GGGGACCACTTTGTACAAGAAAGCTGGGTTCtgctcggagaagaggtgagtg | BP pDONR221 for pFAST-R07 |
| | ACS2-LP<br>ACS2-RP | TTGTCGTTCAATTAAAACCGC<br>AGAATTGACACAGCAAATGGG | genotyping / sequencing |
| **ACS5** | ACS5-LP<br>ACS5-RP | CCAGCTATGTTTCGATCTAATCGAGTCATGGTTAAC<br>GAGGTCAAGCTCTGCTTCAAATGTGTTTGTGTCCA | genotyping |
| **ACS6** | ACS6-LP<br>ACS6-RP | CACTTGGTGAACAATCACACG<br>GCTTGCCTGAATTCAGACAAG | genotyping |
| **ACS7** | pACS7-attB1-F<br>ACS7-attB2-R | GGGGACAAGTTTGTACAAAAAAGCAGGCTTAacgaattgtaaccaacacgcaatgg<br>GGGGACCACTTTGTACAAGAAAGCTGGGTTCaaacctccttcgtcggtccat | BP pDONR221 for pFAST-R07 |
| | ACS7-LP<br>ACS7-RP | AACTTGCTTTGTCCAAGCAAG<br>ATCCTAACGACGCCCTTCTAG | genotyping / sequencing |
| **ACS8** | ACS8-LP | ATAACCAACCCATCTAACCCG | genotyping |

| | | | |
|---|---|---|---|
| | ACS8-RP | GGCTTCTCAACCAGAAAGGTC | |
| **ACS9** | ACS9-LP<br>ACS9-RP | GTTTGAGAAGACACGAGACCG<br>CCTACTTCTTGGGATGGGAAG | genotyping |
| **UBC10** | UBC10-F<br>UBC10-R | CAAGGTGCTGCTATCG<br>ATCTCGGGCACCAAAGG | qRT-PCR |
| | HIS293RV<br>LacZ592RV | GGGACCACCCTTTAAAGAGA<br>ATGCGCTCAGGTCAAATTCAGA | pMW#2 &pMW#3 specific primers. |
| **NLS:RFP** | **nls:RFP-attB1-F**<br><br>**RFP-attB2-R** | GGGGACAAGTTTGTACAAAAAAGCAGGCTTACCCAAGAAGAAGCG CAAAGTGATGGCCTCCTCCGAGGAC<br><br>GGGGACCACTTTGTACAAGAAAGCTGGGTTTAGGCGCCGGTGGAG TG | Positive control for FLIM-FRET assay. |
